# Keratin dynamics govern the establishment of the maternal-fetal interface

**DOI:** 10.1101/2021.04.07.438772

**Authors:** Wallis Nahaboo, Sema Elif Eski, Marjorie Vermeersch, Bechara Saykali, Daniel Monteyne, Thomas M. Magin, Nicole Schwarz, An Zwijsen, David Perez-Morga, Sumeet Pal Singh, Isabelle Migeotte

**Author notes:** Correspondence should be addressed to I.M.

## Abstract

After implantation, the mouse embryo undergoes gastrulation and forms mesoderm and endoderm. Mesoderm participates in embryonic and extra-embryonic tissues including the amnion, yolk sac, chorion and allantois, the umbilical cord precursor.

Extra-embryonic mesoderm is rich in intermediate filaments. Two-photon live imaging of keratin 8-eYFP knock-in embryos allowed recording nucleation and elongation of keratin filaments, which formed apical cables coordinated across multiple cells in amnion, allantois, and blood islands. Embryos lacking all keratins displayed a deflated exocoelomic cavity, a narrow thick amnion, and a short allantois, indicating a hitherto unknown role for keratin filaments in post-implantation extra-embryonic membranes morphogenesis.

Single-cell RNA sequencing of mesoderm cells, microdissected amnion, chorion, and allantois provided an interactive atlas of transcriptomes with germ layer and regional information. Keratin 8^high^ mesenchymal cells in contact with the exocoelom shared a cytoskeleton and adhesion expression profile that might explain the adaptation of extra-embryonic structures to the increasing mechanical pressure.

**GRAPHICAL ABSTRACT:** 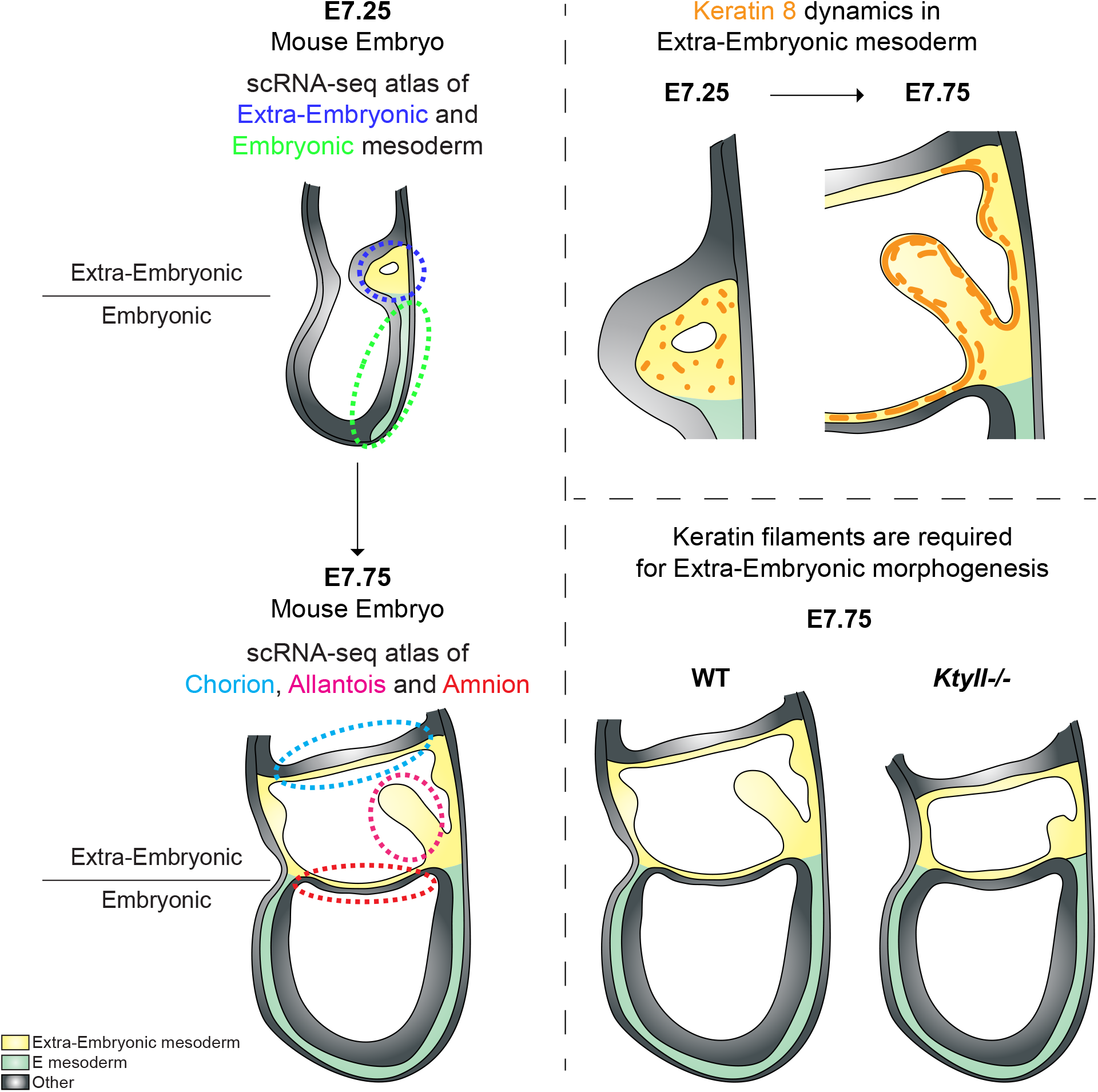

## INTRODUCTION

Placenta was defined as the “apposition or fusion of the fetal membranes to the uterine mucosa for physiological exchange” by H.W. Mossman in 1937. The placenta supports the developing embryo through provision of water, nutrients, and gas exchange ^1^. Early lethality in the mouse is almost always associated with severe placental malformations ^2,3^. Placenta is also a major determinant of post-natal health ^4,5^.

At early stages, mammalian embryos absorb nutrients through endocytosis. Post-gastrulation, embryo development relies on proper morphogenesis of extra-embryonic (ExE) envelopes (Figure 1A). The fetal portion of the placenta comprises trophoblast, mesenchymal and vascular cells, derived from distinct germ layers. Trophoblast (from the Greek words for “feed” and “embryo”) originates from trophectoderm, the external layer of the blastocyst stage embryo, that solely contributes to extra-embryonic structures including ectoplacental cone and ExE ectoderm ^6,7^. Mesenchymal and vascular cells originate from the epiblast, a germ layer derived from the blastocyst inner cell mass. During mouse embryo gastrulation, posterior epiblast cells delaminate at the primitive streak to become endoderm and mesoderm. ExE mesoderm emerges from the most posterior part of the streak and accumulates between ExE ectoderm and visceral endoderm to create the amniochorionic fold, in which fusion of multiple lumens generates the exocoelomic cavity ^8^. Amnion, the innermost ExE tissue, is a thin bilayer formed at embryonic day (E) 7 from epiblast and ExE mesoderm; it separates exocoelomic and amniotic cavities ^9^. In primates and rodents, placenta arises from the fusion of ectoplacental cone, chorion and allantois ^10^. When the amniochorionic fold is fully expanded, chorionic walls formed of ExE ectoderm and mesoderm fuse anteriorly ^8^. Chorion then detaches from amnion and comes in contact with the ectoplacental cone. The allantois (Greek word for “sausage”) is the precursor for the umbilical cord and the vessels of the placental labyrinth ^11^. It appears at E7.5 as an ExE mesoderm bud continuous with the primitive streak and grows through cell migration and division ^12^ in the exocoelom towards the chorion, to which it attaches around E8.5. The fetus becomes dependent on the interaction of maternal and fetal circulations in the placental labyrinth from E10.5 ^4,5^.

**Figure 1:**
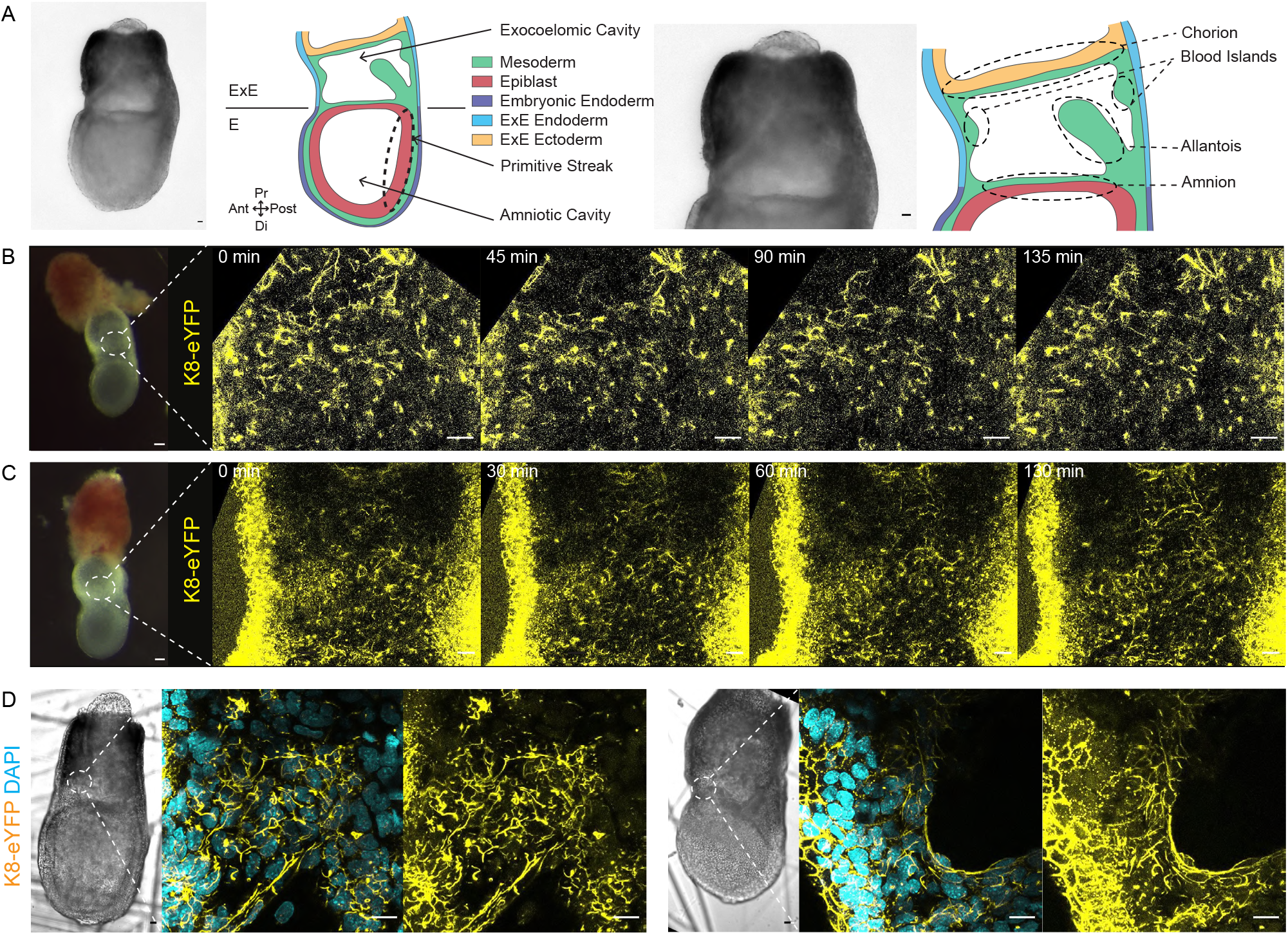
Nucleation of K8 filaments in E7.5 mouse embryo. A. Bright-field picture and scheme of an E7.5 Late Bud (LB) embryo showing germ layers in Embryonic (E) and Extra-Embryonic (ExE) regions (Left). Zoom on ExE region with annotations of ExE structures (Right). B. Bright-field picture of a 0B K8-eYFP embryo (Left) and time series of two-photon live imaging of ExE region (Right) (n=9). C. Bright-field picture of EB K8-eYFP embryo (Left) and time series of two-photon live imaging of ExE region (Right) (n=8). D. Bright-field pictures of K8-eYFP LB embryos (n=30), and high magnification spectral acquisition of areas of ExE mesoderm corresponding to dashed white areas. K8-eYFP and DAPI appear in yellow and cyan, respectively. Scale bars represent 25 μm, except for bright-field pictures in B-C where scale bars show 50 μm.

We recently identified differences in morphology, migration pattern, and gene expression between embryonic and ExE mesoderm ^13^. In particular, intermediate filament components keratins (Krt or K) 8 and 18 are highly abundant in ExE mesoderm. Keratin intermediate filaments (KF) are formed of obligatory heterodimers of type I and type II monomers assembled into tetramers in an antiparallel manner, which associate laterally and longitudinally to generate 8-12 nm filaments ^14^. KF display high tensile strength and toughness, as well as remarkable elasticity, which may allow cells to sustain and recover from large deformations. In epithelial tissues, desmosome-anchored KF form a mechanically resilient transcellular network that is modulated by and protects against physical stress ^15^. In the mouse embryo, K8 and 18 are already detected in a subset of cells at the eight-cell stage ^16^, and it was suggested that they function as asymmetrically inherited factors that specify the first trophectoderm cells ^17^. Establishment of a knock-in K8-eYFP fusion mouse line allowed tracking KF network formation and dynamics in live embryos without functional alteration ^18^. K8-eYFP accumulates as dots at cell borders during the morula stage ^15^. At the blastocyst stage, there is a dense KF network in trophectoderm, and no K8/18 expression in inner cell mass. At E7, immunostaining for K8 marks ExE ectoderm and mesoderm, as well as visceral endoderm ^13^. An important role for KF in mesenchymal tissue was uncovered in the frog embryo, where KF mechanically connect mesendodermal cells and are therefore required for efficient collective migration ^19^.

To adapt to the embryo’s growth and needs, ExE structures need to rapidly expand and change morphology. At E8.5 the U-shaped embryo turns along its dorsoventral axis and becomes enclosed in the amniotic sac in which it matures and moves protected from traumas, infections and toxins. Embryo and amnion are surrounded by the yolk sac that provides nutrition and gas exchange until the placenta is ready to take over. The umbilical cord must accommodate fetal movements and ensure maintenance of blood flow between fetus and mother. Collectively, this requires that amnion, yolk sac and allantois must be both resistant and elastic. As we found high expression of intermediate filament components in ExE mesoderm, we set up to explore keratins’ function in the post-implantation mouse embryo and its supporting tissues.

Because of high and ubiquitous K8 expression in ExE cells, the K8-eYFP line allowed recording both dynamics of KF and morphogenesis of ExE structures through two-photon live microscopy. At late gastrulation stage, live imaging identified KF nucleation and elongation in ExE mesoderm as well as in amnion, allantois, chorion and blood islands. Areas in contact with the rapidly expanding exocoelom, presumably exposed to the highest mechanical constraint, displayed long KF cables spanning several cells. Mutant embryos devoid of all keratins had a collapsed ExE cavity, a short thick amnion and a small allantois, suggesting KF play a major role in shaping ExE envelopes. As profiles of early ExE tissues at cellular resolution were lacking from mouse embryo single-cell atlas, we performed single-cell RNA sequencing analysis (scRNA-seq) of E7.25 mesoderm cells as well as E7.75 microdissected amnion, allantois and chorion, thereby identifying the expression landscape of KF-rich epithelial and mesenchymal cells and providing a detailed atlas of ExE structures.

## RESULTS

### KF nucleate and elongate in ExE mesoderm, forming a network continuous across cells

To record KF dynamics in the post-implantation mouse embryo, we performed two-photon static and live imaging of Early Streak (ES, E6.25) to Late Bud/Neural Plate (LB/NP, E7.75) stages ^20^ *ex vivo* cultured K8-eYFP embryos ^18^ (Figure 1A, Sup1A). As expected, eYFP was present in visceral endoderm, ExE ectoderm, and ExE mesoderm (Sup1B, C). 3D-reconstruction of LB embryos stained for F-actin showed that, similar to E7.25 ExE mesoderm ^13^, ExE mesenchyme in amnion, chorion and allantois had low F-actin, compared to embryonic mesoderm (Sup1D and Video 1). From Late Streak (LS)/Zero Bud (0B) stages, eYFP positive dots (5,34±0,62 μm², n=24 from 6 embryos) could be detected specifically in ExE, but not in embryonic, mesoderm cells (Figure 1B, Sup1C and Video 2). Dots subsequently elongated into filamentous particles up to 25 μm (24,76±2,47 μm, n=19 from 8 embryos) in length, and organized a reticular web (Figure 1C and Video 3). Higher resolution imaging of Early Bud (EB) to NP fixed samples showed that K8 containing filaments could form linear structures up to 75 μm in length (65±11,92 μm, n=11 from 6 embryos) continuous across multiple cells, in particular in the walls of the expanding ExE envelop as well as in the developing amnion (Figure 1D and Video 4).

### Stretchable KF-rich cables connect amnion, exocoelomic wall, and chorion

Live imaging of K8-eYFP embryos recorded the global morphogenetic events that shape ExE structures (Figure 2). The amnion displayed small excrescences perpendicular to its ExE surface. We noticed the formation of stable (> 3 hours) K8 containing “cables”, mostly in the anterior part of the embryo, bridging the mesoderm-derived part of the amnion and the wall of the exocoelomic cavity (Figure 2A and Video 5). Those cables stretched progressively (initial length 52,61±15,37 μm and width 9,69±5,8 μm, final length 130,5±48,98 μm and width 5±3,41 μm, n=5) and finally snapped as the cavity grew; they likely represent remnants of connections between amnion mesoderm and blood islands that break upon amnion closure. Cables linking amnion and wall were not detected in fixed samples, suggesting they might be unstable and lost upon fixation. In a subset of embryos (n=6 with the adequate stage and orientation), a larger KF-rich structure was visible in the anterior region (Figure 2B and Video 6), corresponding to the chorion portion of the amniochorionic fold. It grew longer and thinner before rupturing at the Anterior Separation Point (Sup1A) ^8^, allowing full expansion of the exocoelom and positioning of the chorion close to the ectoplacental cone. Higher resolution imaging of fixed samples highlighted a dense KF network on the amnion ExE side and within the retracting chorion (Figure 2E, E’).

**Figure 2:**
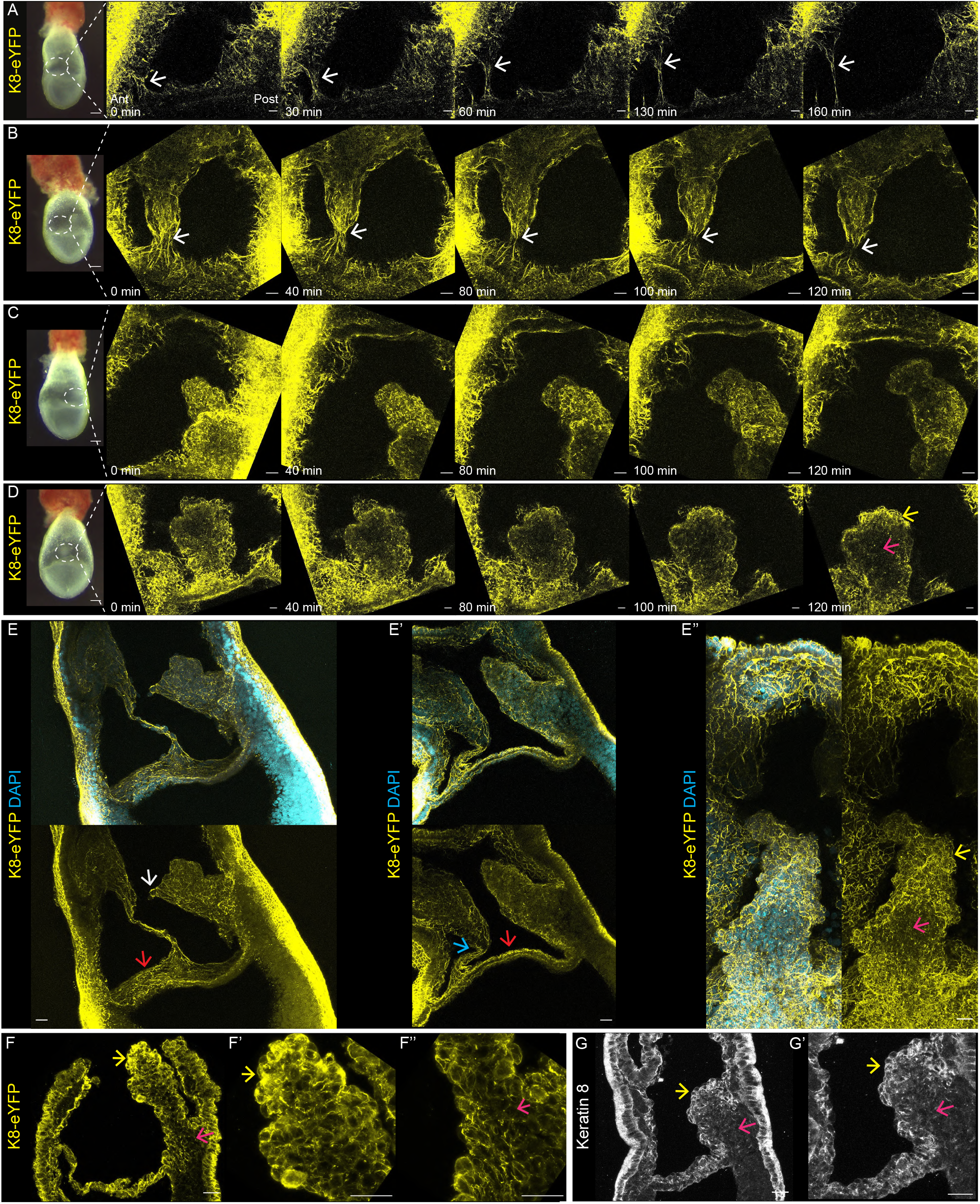
K8 filaments extend in ExE region. A-D. Bright-field pictures of K8-eYFP (Left) and Z-projections of time series (Right) of live LB K8-eYFP embryos imaged using two-photon laser with time intervals between 20 and 30 min focusing on regions corresponding to dashed white areas. A. Lateral view showing extension of K8 containing cables (white arrows) attached to amnion and ExE wall (n=10). B. Anterior view showing chorion retraction at the Anterior Separation Point (white arrows) (n=6). C. Lateral and D. Posterior views showing allantois growth over time (n=12 and 3). Allantois cortex and mesenchyme are indicated with yellow and magenta arrows, respectively. E. Spectral images at high magnification of ExE region from fixed K8-eYFP LB/NP embryos (n=20). E: lateral view where white and red arrows point to allantois edge and amnion. E’: lateral view where blue and red arrows show Anterior Separation Point and amnion. E’’: posterior view focused on allantois where yellow and magenta arrows indicate allantois cortex and mesenchyme. K8-eYFP and DAPI appear in yellow and cyan. F. Immunostaining on sagittal section (anterior to the left) of a LB K8-eYFP embryo (n=20) with anti-eGFP antibody (yellow) focusing on ExE region (F); higher magnification images of allantois edge (F’, yellow arrow) and base (F’’, magenta arrow). G-G’. Immunostaining on sagittal section of a LB wild-type embryo (n=32) with anti-Krt8 antibody (white) (G); higher magnification acquisition centred on allantois (G’) where yellow and magenta arrows indicate allantois cortex and mesenchyme. Scale bars represent 25 μm, except for bright-field pictures in A-D where scale bars show 100 μm.

### KF form a shell around the allantois

Allantois growth towards the chorion was recorded on the posterior side of the embryo. The allantois had an irregular shape and expanded through formation of smooth blebs and bulges. The directionality of extension appeared stable. K8-eYFP uncovered a striking difference between the external layer of the allantois, which has been referred to as “mesothelium” ^21^ and is called “cortex” in this study, and the inner allantois cells. The cortex displayed a rich KF network, while the inner allantois cells had a lower KF content (Figure 2C, D and Video 7). 3D reconstruction from whole-mount imaging of fixed samples illustrated the reticular network covering the entire allantois (Figure 2E-E” and Videos 8a, b). Interestingly, the allantois cortex displayed KF-rich cell blebs (as described in ^21^), some of which were directed towards reciprocal bubbles in chorion mesenchyme (Figure 2E). Comparing sections from K8-eYFP embryos (Figure 2F-F”) and wild-type embryo sections immunostained for K8 (Figure 2G-G’) confirmed that K8-eYFP fusion protein recapitulates native K8 expression pattern.

### ExE cell populations have distinct morphological characteristics

Scanning electron microscopy (SEM) of LB/NP embryos provided valuable insights on ExE tissue and cell morphology (Figure 3A). Blood islands were distinguished on the cavity wall, notably through the emergence of tubular structures (Figure 3B). The allantois had a “cotton candy” appearance with multiple blebs (Figure 3C-C’) ^21^. Opening the allantois confirmed distinct cellular shapes between cortex and mesenchyme (Figure 3D). Allantois cortex consisted of a cohesive layer of apically concave stretched cells with short villi (Figure 3C-C’). In contrast, inside cells were rounder; they displayed long entangled filopodia mostly concentrated at cell-cell junctions and a network of twisty, branched filaments (Figure 3E-E’). In the amnion, SEM highlighted the difference between flat epiblast-derived cells on the embryonic side (Figure 3F) and mesoderm-derived cells forming a hilly landscape on the ExE side (Figure 3G-G’). Transmission electron microscopy (TEM) uncovered subcellular details, such as filaments and adhesion complexes. In blastocyst trophectoderm, KF nucleation sites were shown to co-localize with nascent desmosomes ^15^. In the allantois (Figure 3H-J), we found tight and adherent junctions in inner cells (Figure 3I-I’), whereas external cells were predominantly connected by adherent junctions (Figure 3J). Microfilaments adjacent to adherent junctions were detected at higher magnification (Figure 3J’). In the amnion (Figure 3K-M), tight and adherent junctions were found in epiblast (Figure 3L) and mesoderm (Figure 3M-M’)-derived cells. In addition, atypical, possibly immature, desmosome-like junctions were observed (Figure 3L’). Filaments bundles and desmosomes were abundant in chorion (Figure 3N-N’). Interestingly, TEM unveiled the presence of primary cilia specifically in cuboidal cells on the posterior side of the allantois base, where high and localized Hedgehog signaling was previously identified through a *Ptc1*:lacZ reporter ^22^ (Figure 3O-O’). Those cells displayed nascent desmosomes (Figure 3O’’), compatible with their lower permeability to dextran ^21^.

**Figure 3:**
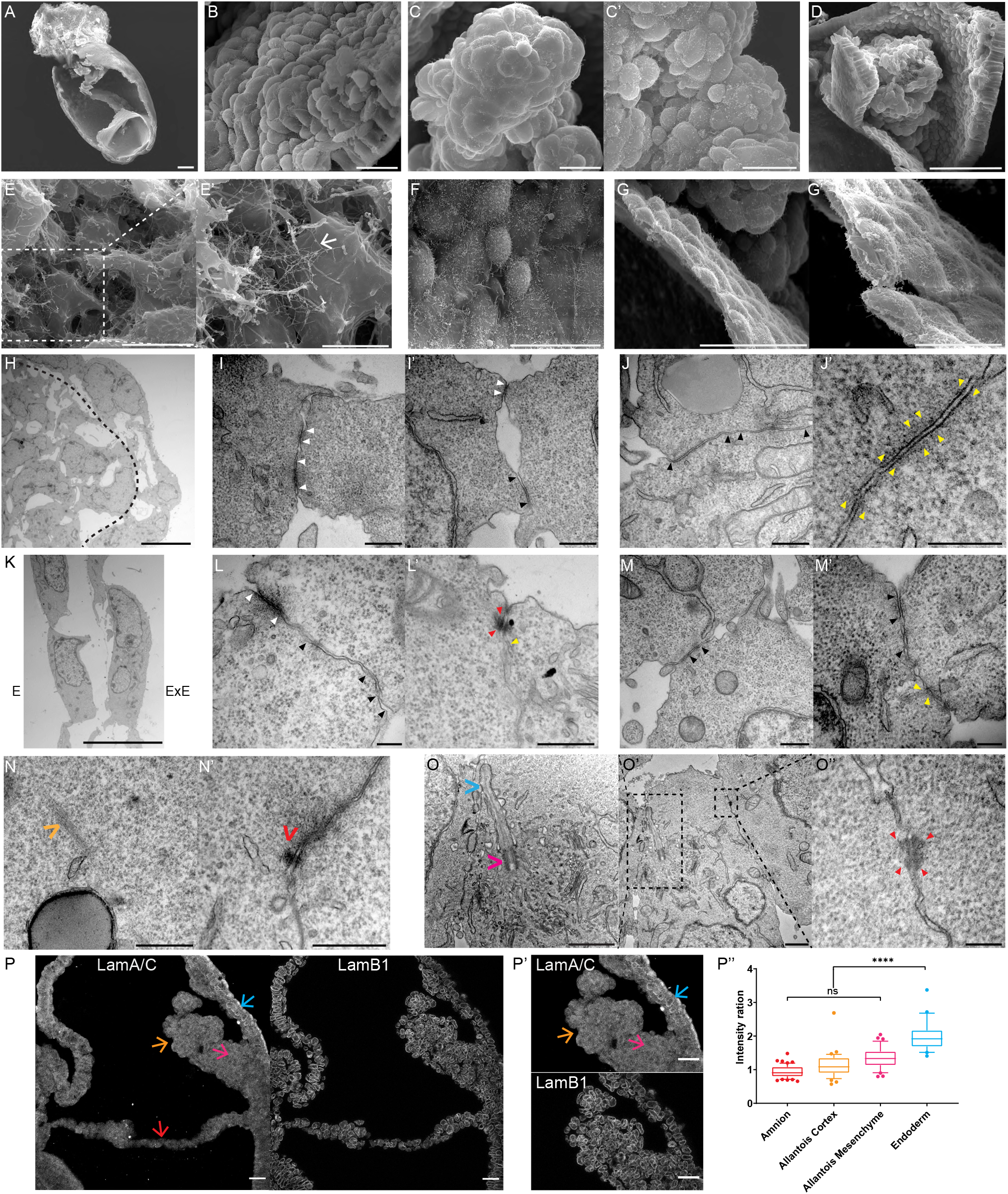
Electron microscopy of the ExE region at LB/NP stage. A-H. SEM images of an opened embryo (A), the ExE cavity wall (B), the allantois surface (C-C’), an opened allantois (D) and its zooms (E-E’) where filaments are indicated by white arrows, the epiblast-derived side of the amnion (F), the mesoderm-derived side of the amnion (G) and its zoom (G’). H-O. TEM pictures in allan-tois (H-J), amnion (K-M), chorion (N) and allantois base (O). In H, allantois mesenchyme (Left) and cortex (Right) are separated by a black dashed line. I-J. Junctions in internal (I-I’) and external (J-J’) allantois cells. In K, the embryonic side of the amnion is on the left, and the ExE side on the right. L-M. Junctions in the epiblast-derived side (L-L’) and the mesoderm-derived side (M-M’) of the amnion. White and black dashed rectangles represent zoomed regions. Arrowheads point towards tight junctions (white), adherent junctions (black), desmosome-like junctions (red), and filaments (yellow). In chorion, the orange arrow indicates intermediate filaments (N) and the red arrow a desmosome (N’). In O, axonem (cyan arrow) and basal body (magenta arrow) are annotated. Scale bars represent 100 μm (A, D), 50 μm (G), 25 μm (B, C, E, F), 20 μm (G’) 10 μm (H, K, P), 5 μm (E’), 1 μm (O, O’), 500 nm (I, J, L’, M, N), and 200 nm (J’, L, M’, O’’). P. Sagittal sections of a NP embryo immunostained for LamA/C and LamB1 (P-P’). Quantification (P’’) of the ratio of LamA/C over LamB1 signal intensity in amnion (red), allantois cortex (yellow), allantois mesenchyme (magenta), and visceral endoderm (cyan) (P’) (n=6 embryos). Boxes extend from the 25th to 75th percentiles, and whiskers are drawn down to the 10th percentile and up to the 90th. Points below and above the whiskers are indicated as individual points. Scale bars represent 25 μm. P-values were calculated using the Mann-Whitney-Wilcoxon test.

### The nuclear membrane composition of ExE cells suggests an elastic behavior

The shape and membrane composition of the nucleus are correlated to the physical properties of the cell ^23^. A high Lamin A/C over Lamin B1 ratio reflects a viscous environment whereas a low ratio is associated to an elastic environment ^24,25^. In the EB/NP mouse embryo, the only cells with high Lamin A/C were in visceral endoderm, suggesting that most tissues, in particular ExE mesoderm-derived structures, have low stiffness and high deformability (Figure 3P-P’’).

### Embryos lacking keratin display major defects in ExE structures morphogenesis

Due to redundancy among keratins, it is difficult to interpret phenotypes resulting from knocking-out individual genes. Deletion of *K8* caused an embryonic lethal phenotype at E12.5 associated with a placental defect ^26^. Combined deletions of *K18/K19* ^27^ or *K8/K19* ^28^ caused fragility of giant trophoblast cells and extensive hemorrhages, which led to death at E10. As KF assembly relies on obligate heterodimerization of members of each of the two families, deletion of the whole keratin family cluster II plus the flanking type I *K18* (*KtyII*^−/−^) allowed to fully overcome redundancy and led to growth retardation from E8.5. Mutant embryos arrested around E9.5 due to defective energy metabolism resulting from abnormal intracellular distribution of glucose transporters in the yolk sac ^29^. To explore the function of keratin in morphogenesis of ExE mesoderm-derived structures, we investigated earlier stages of development.

*KtyII*^−/−^ embryos displayed specific phenotypes as early as E7.5 (Figure 4A): their ExE region was collapsed, with a narrow cavity and scrambled tissue architecture. Immunostaining for K8 at E8.5 confirmed absence of the protein (Figure 4B). Mutant embryos arrested around E9, as previously described (Figure 4C) ^29^. The severity of the phenotype was variable at earlier stages. We quantified the size of ExE structures (Figure 4D) and normalized values by the length of the embryonic region, which was unaffected in mutants (Figure 4E). In EB/NP *KtyII*^−/−^ mutants, the exocoelomic cavity expansion was impaired (Figure 4F), the allantois was smaller (Fig4G), and the amnion was shorter and thicker (Figure 4H, I). The phenotype of E7.5 embryos lacking keratin argues for a predominant function of KF in ExE mesenchymal tissues, in addition to epithelia. Aligned KF in the exocoelomic wall and amnion are likely to support elasticity, while KF might preserve tissue rigidity of the allantois cortex during elongation.

**Figure 4:**
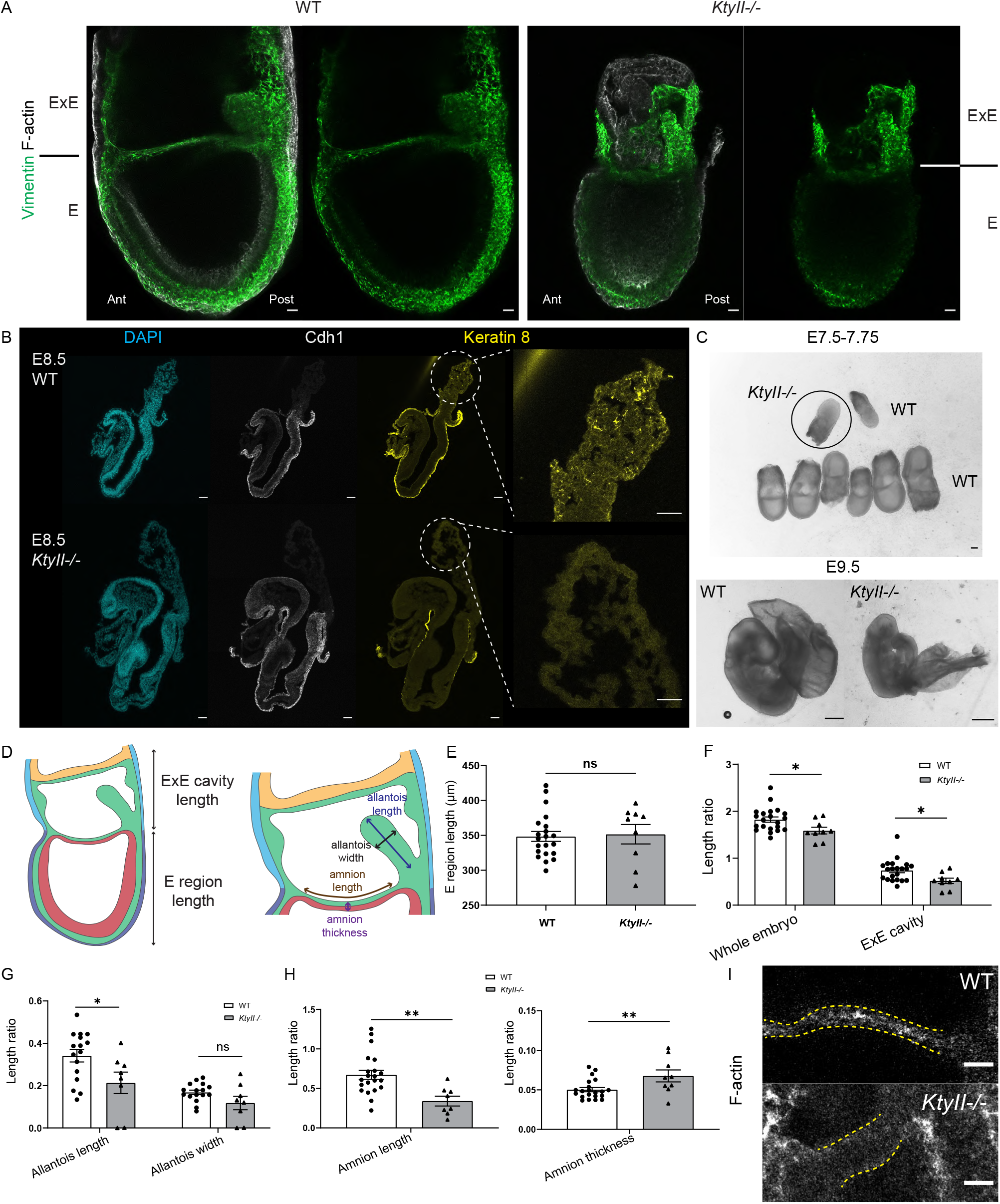
Loss of keratin results in defective ExE region morphogenesis. A. Z-projections of whole-mount wild-type (WT, Left) and KtyII−/− (Right) embryos at LB stage stained for vimentin (green) and F-actin (Phalloidin, white). B. Sagittal sections of wild-type (WT, n=6) and KtyII−/− (n=2) E8.5 embryos stained for nuclei (DAPI, cyan), Cdh1 (white) and Krt8 (yellow). There is some non-specific staining on the section edges but none in allantois (zoom). C. Bright-field images of E7.5-7.75 LB litter (top) and E9.5 (bottom) WT (left) and KtyII−/− (right) embryos. Scale bars represent 100 μm. E. Scheme for measurements. E-H. Quantification of embryo morphology in wild-type (white boxes/ black dots, n=22 including 16 in which allantois could be measured) and KtyII−/− embryos (grey boxes/ black triangles, n=9 including 8 in which allantois could be measured) at LB stage. (E) Embryonic region length. Ratios of whole embryo and ExE cavity lengths (F), allantois length and width (G) and amnion length and thickness (H), over embryonic region length. P-values were calculated using the Mann-Whitney-Wilcoxon test. Error bars represent SD. I. Z-projections of wild-type (Top) and KtyII−/− (Bottom) embryos at LB stage stained for F-actin (Phalloidin, white). Amnion is delimited by yellow dashed lines. Scale bars represent 25 μm.

### Temporal and spatial dynamics of the transcriptional landscape in K8^high^ cells

Despite the deformations associated with keratin loss, most mutant embryos successfully completed turning and displayed allantois elongation, indicating that other factors might compensate for the lack of KF in order to allow amnion rapid stretch and allantois growth. To get a global view of ExE mesenchymal cells diversity in time and space, we turned to scRNA-seq. An interactive atlas was created to facilitate usage of the clustered data for the community. We previously performed bulk mRNA analysis of Mid/Late Streak (M/LS) stage mesoderm cells recovered from *Brachyury*-Cre; mTmG transgenic embryos, in which Cre-recombination in cells expressing *Brachyury* activates a membrane GFP reporter while non-recombined cells express membrane Tomato (Figure 5A). Here M/LS Embryos from the same genetic background were manually cut at the embryonic-ExE boundary, GFP+ cells from each population were sorted by flow cytometry (Sup2A), and single cells were sequenced. RNA velocity, which reveals the stream from immature to mature cell state ^30^, assisted in cluster identification.

**Figure 5:**
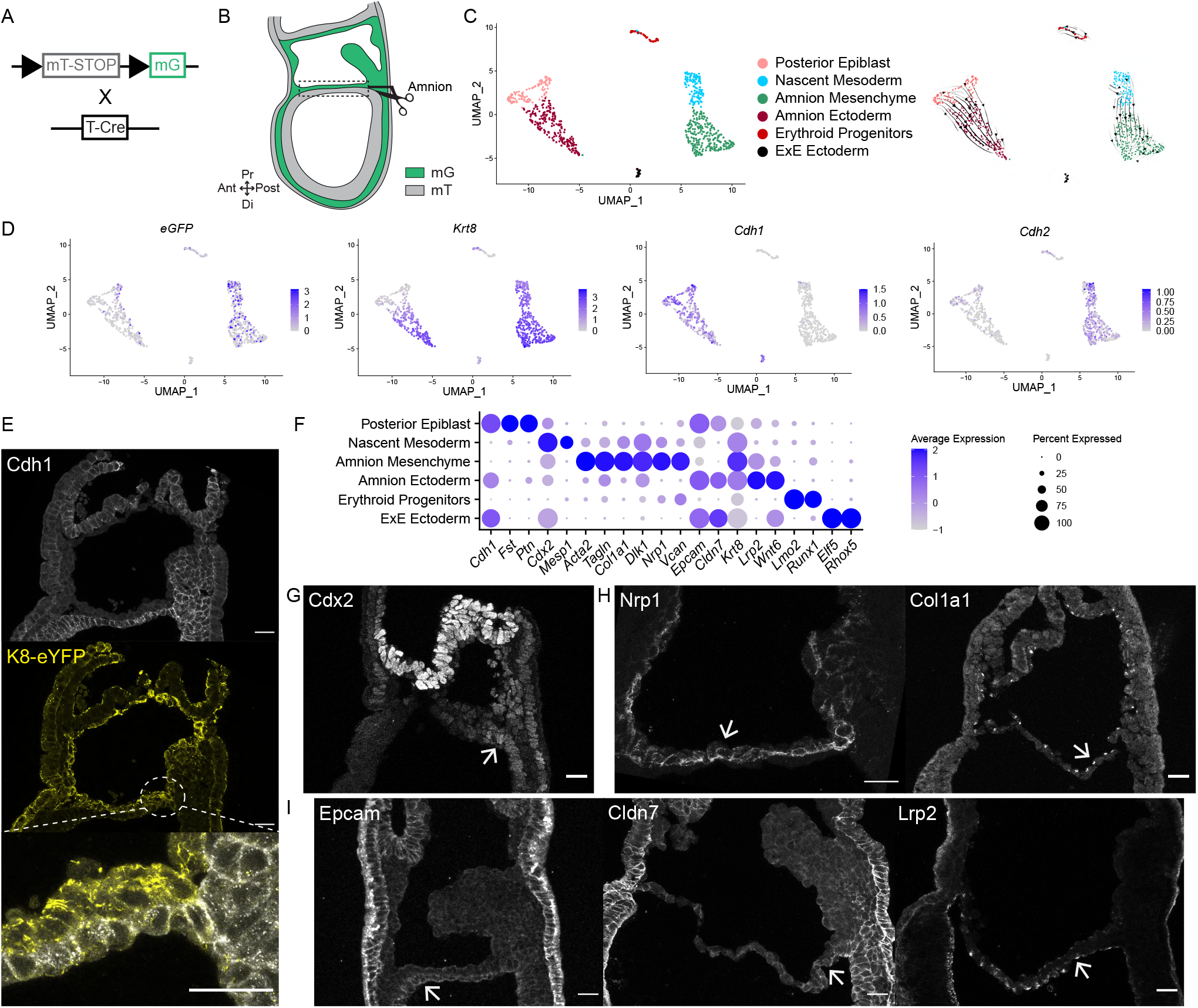
Amnion single cell transcriptome. A. In Brachyury (T)-Cre; mTmG embryos, mesoderm-derived cells express membrane eGFP (mG, green) and the other cells membrane tdTomato (mT, grey). B. Scheme of amnion isolation. C. Uniform Manifold Approximation and Projection (UMAP) (Left, color-coded) and RNA velocity (Right, black arrows show streaming directions from unspliced to spliced RNA) of amnion cells where 6 unsupervised clusters were identified. D. UMAP of eGFP, Krt-8, Cdh1 and Cdh2. E. EB K8-eYFP embryo stained for eYFP (yellow) and Cdh1 (white) (n=7). Dashed white area zoom on an amnion region which highlights the two cell layers (Bottom). F. Dot plot of specific gene expression in amnion clusters. G. Immunostaining for Cdx2 (n=20). H. Immunostaining for Nrp1 (n=8) and Col1a1 (n=7), primarily expressed in Amnion Mesenchyme. I. Immunostaining for Epcam (n=12), Cldn7 (n=7), and Lrp2 (n=6), primarily expressed in Amnion Ectoderm. White arrows point toward signal in amnion. In (E) and (G-I), sagittal sections with anterior to the left. Scale bars represent 25 μm.

Embryonic mesoderm cells were relatively homogeneous, as expected at this stage of gastrulation. Based on gene expression and RNA velocity streams, we defined six clusters (Sup2B, C). Five of those were adjacent: Primitive Streak, Nascent Mesoderm, Lateral Plate/Paraxial Mesoderm, Cranial/Heart Mesoderm, and ExE Mesoderm. A small independent cluster corresponded to Node precursors. Among the genes previously identified as enriched in embryonic mesoderm, single cell analysis showed particularly interesting profiles for guidance molecules (Sup2D). *Epha1* and *Ntn1*, for example, were found predominantly in the streak and node precursors, suggesting a role in transitions between epithelial and mesenchymal stages, while *EphA4* was expressed in Nascent and Paraxial/Lateral Mesoderm, but not in Anterior and ExE mesoderm, compatible with its later function in somitogenesis ^31^. For ExE mesoderm cells, we distinguished three clusters (Sup2E-G): Primitive Streak, Endothelial/Erythroid Progenitors, and ExE mesenchyme. Apart from blood and vessels precursors, it was difficult to predict cell fate based on expression profiles. Indeed, cells from different origins can become undistinguishable once they either acquire a particular fate, such as gut endoderm which derives from both definitive and visceral endoderm ^32^, or migrate to a particular region, as illustrated by high K8 expression in all ExE layers.

To better understand cell differentiation trajectories, we characterized ExE populations at later stages and combined spatial and germ layer information. We manually isolated amnion, allantois, and chorion from *T*-Cre; mTmG embryos dissected at LB/NP stages (Figure 5, 6, 7). All cells were sequenced to obtain a comprehensive atlas of each tissue. In addition to previously described expression profiles, each cluster was validated by immunostaining for at least one specific marker.

**Figure 6:**
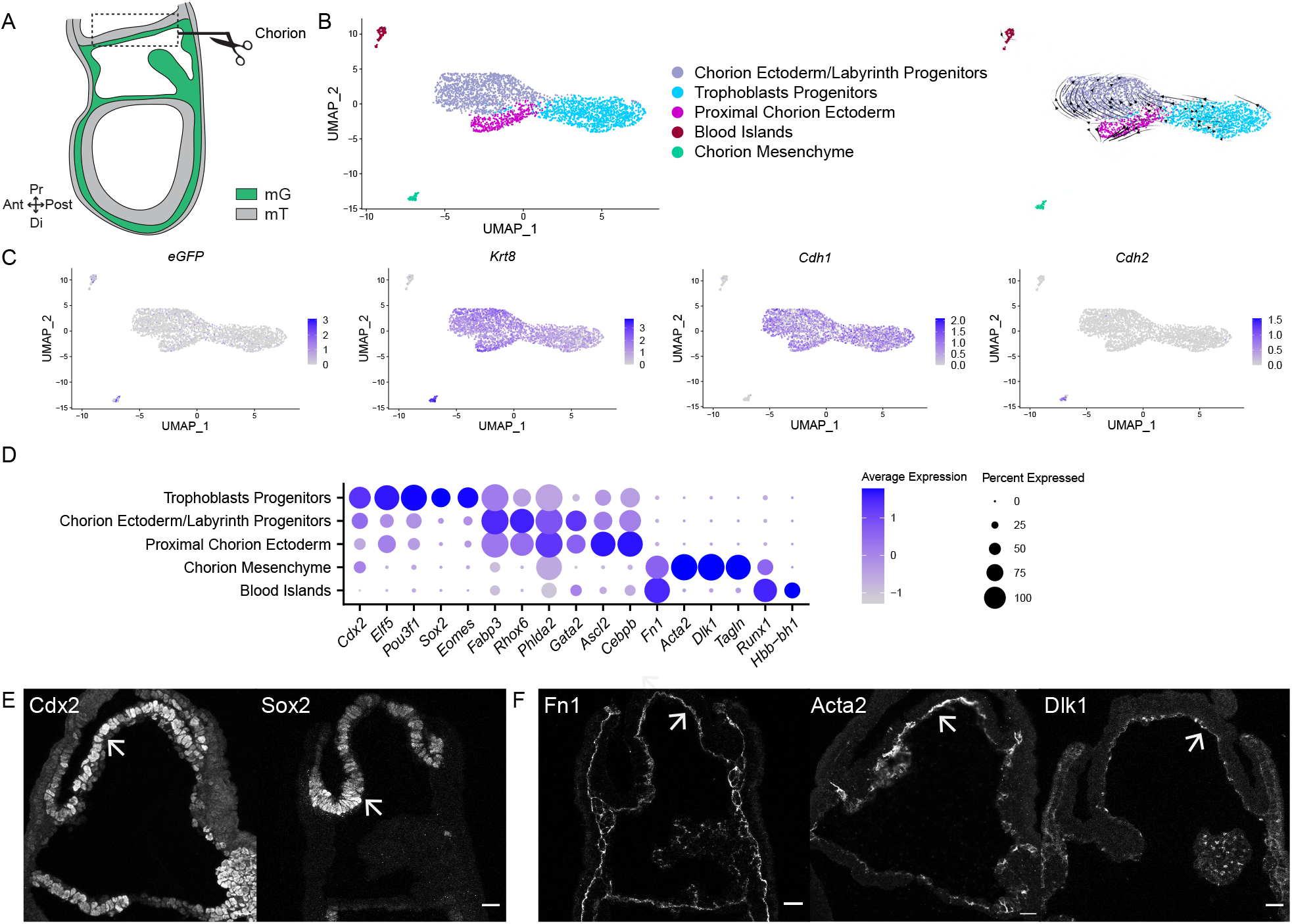
Chorion single cell transcriptome. A. Scheme of chorion isolation where mesoderm is in green (mG) and the other cells are in grey (mT). B. UMAP (Left) and RNA velocity (Right) of chorion cells where 5 unsupervised clusters were identified. C. UMAP of eGFP, Krt-8, Cdh1 and Cdh2. D. Dot plot of specific gene expression in chorion clusters. E. Immunostaining for Cdx2 (n=20), Sox2 (n=10), primarily expressed in Trophoblasts Progenitors. F. Immunostaining for Fn1 (n=22), Acta2 (n=12), and Dlk1 (n=8), primarily expressed in Chorion Mesenchyme. White arrows point toward signal in chorion. Sagittal sections with anterior to the left. Scale bars represent 25 μm.

**Figure 7:**
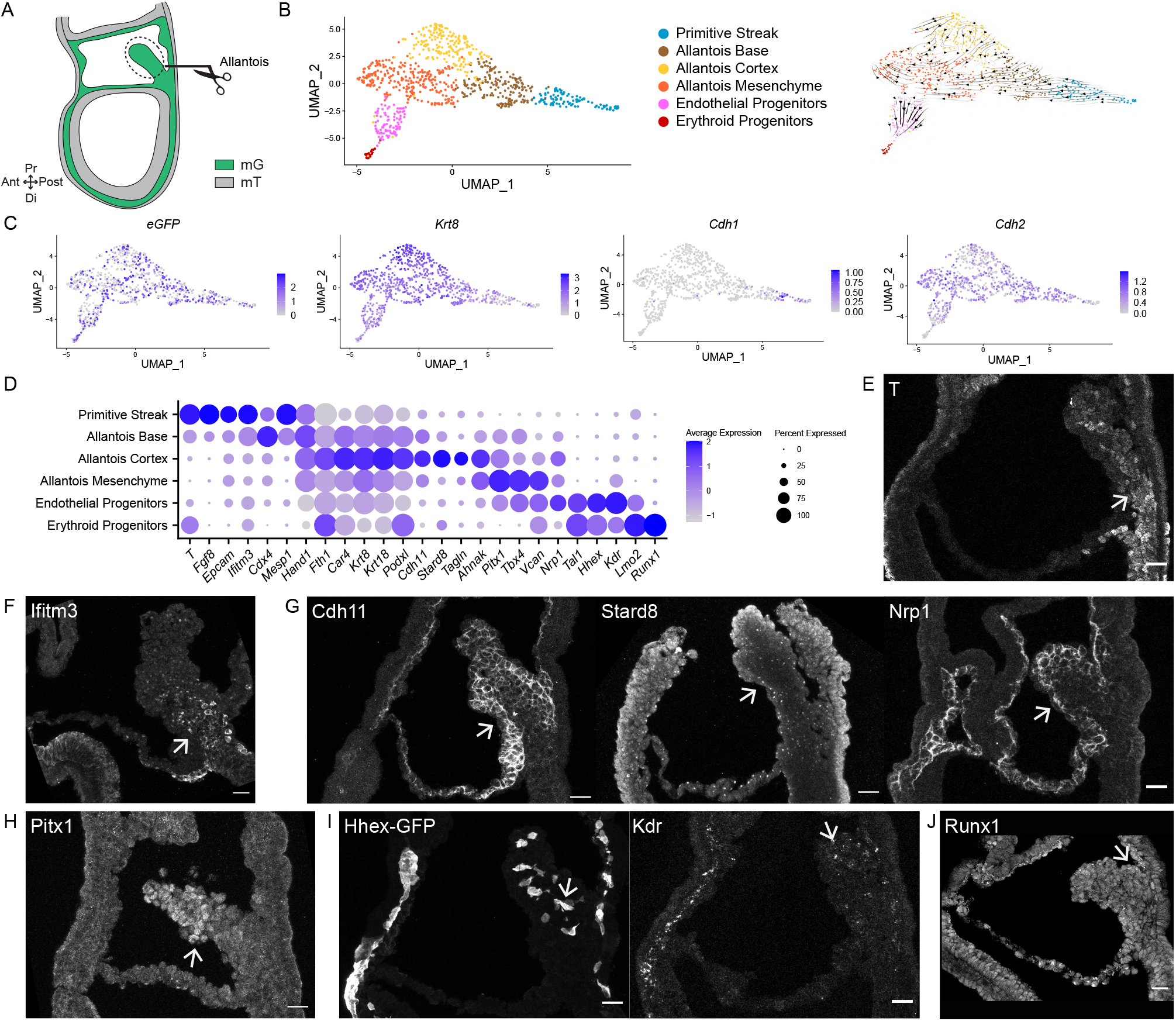
Allantois single cell transcriptome. A. Scheme of allantois isolation where mesoderm is in green (mG) and the other cells are in grey (mT). B. UMAP (Left) and RNA velocity (Right) of allantois cells where 6 unsupervised clusters were identified. C. UMAP of eGFP, Krt-8, Cdh1, and Cdh2. D. Dot plot of specific gene expression in allantois clusters. E. Immunostaining for T (n=5) (Primitive Streak and Allantois Base). F. Immunostaining for Ifitm3 (n=7) (Allantois Base). G. Immunostaining for Cdh11 (n=9), Stard8 (n=4) and Nrp1 (n=8) (Allantois Cortex). H. Immunostaining for Pitx1 (n=4) (Allantois Mesenchyme). I. Hhex-GFP (n=5) and immunostaining for Kdr (n=8) (Endothelial Progenitors). White arrows point toward signal in allantois. Sagittal sections with anterior to the left. Scale bars represent 25 μm.

Amnion cells segregated in 5 clusters regrouped in 2 major populations (Fig5C). One had high *eGFP*, *Krt8* and *Cdh2* and comprised Nascent Mesoderm (*Cdx2, Mesp1*) and Amnion Mesenchyme (*Acta2, Tagln, Col1a1, Dlk1, Nrp1, Vcan*). The low *eGFP* population represented Posterior Epiblast (*Cdh1, Fst, Pst*) and Amnion Ectoderm (*Epcam*, *Cldn7, Krt8, Lrp2, Wnt6*) (Figure 5D-F, Sup3A, B). Amnion also comprised a population of Erythroid Progenitors (*Lmo2/Runx1*) (Sup3C). A small ExE ectoderm cluster (*Elf5, Rhox5*) was likely formed by non-amnion cells that came along during dissection. Immunostaining showed a distinct pattern for K8 in E-cadherin (Cdh1) positive epiblast-derived cells, where it appeared as apical dots, compared to mesoderm-derived cells where K8 was more abundant and formed filaments (Figure 5E). Cdx2 marked nascent mesoderm exiting the streak to become amnion mesoderm, with a posterior to anterior gradient (Figure 5G). Amnion mesoderm showed high expression of Col1a1 and Nrp1, similar to exocoelomic cavity walls and chorion mesenchyme (Figure 5H, Sup3A). Amnion ectoderm, compared to epiblast, had similar Epcam, lower E-cadherin and Claudin7, and higher K8 (Figure 5E, F and I, Sup3B). Specific markers included *Wnt6* and *Lrp2* (Figure 5F). Immunostaining for Lrp2 showed a punctate pattern in amnion ectoderm (Figure 5I, Sup3B), compatible with its function in endocytosis.

In chorion, we defined 5 clusters (Figure 6B-D): Chorion Ectoderm/Labyrinth Progenitors, Trophoblast Progenitors, Proximal Chorion Ectoderm, Blood Islands, and Chorion Mesenchyme. *eGFP*+ cells formed a discrete *Cdh2* positive mesenchymal cluster, and additional cells were scattered among the *Cdh1* positive population, compatible with *Brachyury* expression in ExE ectoderm ^12^ (Figure 6C). *Krt8* was present in both epithelial and mesenchymal cells (Figure 6C). *Cdx2* and *Elf5* were present in most cells, except mesenchyme and blood islands (Fig6D and E, Sup4A). RNA Velocity (Figure 6B) suggested two streams. One started from trophoblast progenitors (*Sox2* (Figure 5E, Sup4A*), Eomes*) and one from a chorion ectoderm population (*Rhox6* ^33^, *Cebpb* ^34^) comprising labyrinth progenitors (*Gcm1*, found on the tip of villi in the chorion-derived component of labyrinth ^34^ (Sup4B)). Both converged on a population displaying a gene profile close to ectoplacental cone (*Ascl2,* an imprinted gene expressed in trophectoderm then mostly restricted to ectoplacental cone ^35^, and *Dlx3,* predominantly detected in ectoplacental cone ^34^ and required for labyrinth morphogenesis ^36^ (Sup4B)). *Dlx3* and *Gcm1* were essentially mutually exclusive (Sup4B), consistent with Dlx3 inhibition of Gcm1 transcriptional activity ^37^. *Gcm1+* cells already had a distinctive transcriptomic profile at that early stage (Sup4B). Mesoderm-derived cells (*Fn1*) were subdivided according to their prospective endothelial (*Dlk1, Acta2*) or erythroid (*Hbb-bh1*) identities (Figure 6D and F, Sup4C). Imprinted genes were found in all chorion populations independent of their germ layer of origin. There were also multiple genes involved in transmembrane transport (*Slc38a4* ^38^), endocytosis (*Vamp8*) and metal metabolism (*Mt1, Mt2, Fthl17a*), reflecting chorion’s prospective role in embryo nutrition.

Allantois, as expected, was essentially composed of mesenchymal (*eGFP, Cdh2*) cells with high *Krt8* expression (Figure 7B, C). Based on known gene expression profiles and RNA velocity streams, 6 clusters were individualized (Figure 7B, D): Primitive Streak (*T* (Fig7E, Sup5A ^12^), *Fgf8*), Allantois Base (*Cdx4*, *T*, *Ifitm3* (Figure 7E, F), Allantois Cortex (*Krt8* and *18*, *Stard8*, *Cdh11*, *Nrp1* (Figure 7G, Sup5B)), Allantois Mesenchyme (*Tbx4, Vcan*, *Pitx1* (Figure 7H, Sup5C)), Endothelial Progenitors (*Tal1*, *Hhex*, *Kdr* (Figure 7I, Sup5D)) and Erythroid Progenitors (*Runx1* (Fig7J), *Lmo2*). Using a polygenic score, we defined a small subpopulation of Primordial Germ Cells (PGC) within the Primitive Streak cluster (Sup5E), providing a snapshot of the PGC expression profile while in the allantois and highlighting the need to combine multiple markers, as none appeared PGC-specific. Allantois mesenchyme had high *Versican (Vcan)* expression, compatible with the importance of proteoglycans in protecting blood vessels in the umbilical cord ^39^. Vasculogenesis has been described to start at the distal end of the allantois and to progress proximally ^12^. We were able to identify early expression profiles of specific endothelial populations, such as potential artery progenitors expressing *Dll4* ^40^ (Sup5F). Although KF in allantois cortex appeared continuous across several cells, we could not visualize mature desmosomes by electron microscopy; scRNA-seq detected desmosome components (such as *Dsp*, *Dsc2* or *Pkp4*) in allantois cells, without cluster specificity. Compatible with TEM data, tight junction constituents *Cldn12*, *Tjp1* and *2* were present in cells from multiple clusters. Among cadherins, *Cdh11* stood out as being more abundant in external allantois cells. Similar to amnion and chorion mesoderm, allantois cortex expressed *Acta2* and *Tagln*. It was also enriched for transcripts of genes associated with apical identity, such as *Podxl,* endosomal transport (*Stard8*, also called *Dlc3* ^41^), as well as transmembrane transport of nutrients (*Scl2a1),* ions *(Fth1)* and gases *(Car4*), suggesting that outer allantois cells start acquiring functions that will be essential for the functionality of the labyrinth even prior to attachment to the chorion.

## DISCUSSION

KF have a higher elasticity than actin and microtubules filaments; they can stretch several times their initial length before reaching the yield point. They play a major role in cell resistance to external mechanical stress, as illustrated by the lower elasticity and higher deformability of keratinocytes lacking keratin ^42,43^. Here, live imaging of mouse embryos with a knock-in K8-eYFP fusion allele identified KF containing cables in all tissues facing the exocoelomic cavity (yolk sac, amnion and chorion mesenchyme, allantois cortex), reminding of the supracellular actomyosin network described in wound healing ^44^, *Drosophila* germ band expansion ^45^, and mouse embryo neural tube closure ^46^. Embryos devoid of keratin had a small exocoelom, a short and thick amnion, as well as defective allantois elongation. In keratinocytes, keratin loss did not obviously affect actin and microtubular networks ^43^. We previously showed that in E7.25 embryos actin filaments were less abundant in ExE, compared to embryonic, mesoderm. At later stages, we did not detect supracellular F-actin cables around the exocoelomic cavity or at amnion borders. Altogether, these observations suggest that the KF network is a prominent cytoskeletal determinant of cell resistance to mechanical challenges during ExE membranes formation. Similarly, human fetal mesenchyme and amniotic epithelium co-express keratin and vimentin ^47,48^. Cytoskeletal networks in amniochorion participate to fetal membranes’ capacity to withstand increase in intrauterine pressure during contractions, and recover from mechanical trauma such as amniocentesis ^49^.

Smooth muscle alpha-2 actin (*Acta2*), transgelin (*Tagln,* also called *SM22alpha*), and cadherin-11 were co-expressed with K8 in amnion and chorion mesenchyme as well as in allantois cortex. *Acta2*, *Tagln* and *Cdh11* are TGFβ-inducible genes, compatible with high BMP signaling in the ExE region. Acta2 is a marker for myofibroblasts and smooth muscle cells. It plays an essential role in vascular contractility, and *ACTA2* mutations in human cause a variety of vascular diseases ^50^. Transgelin binds directly to actin filaments and was shown *in vitro* to cause actin gelation due to conversion of loose filaments into a tangle meshwork ^51^. Cadherin-11 is a type II classical cadherin predominantly expressed in mesenchymal tissues ^52^. During embryonic development, it is required for neural crest survival and migration ^53^. Its expression in fibroblasts is correlated with mechanical stress; switching from cadherin-2 to 11 is associated with transition from low contractile migratory fibroblast to highly contractile myofibroblast ^54^. In frog mesendoderm, local forces applied to C-cadherin result in recruitment of KF to cadherin complexes via plakoglobin ^55^. It is therefore conceivable that cadherin-11 might act as a mechanosensor and mediate reorganization of KF at sites of higher tension. Lineage analysis showed that amnion, chorion and allantois mesenchymal cells derive from common mixed fate progenitors in the posterior epiblast ^9^. Single cell transcriptome of MS/LS ExE mesoderm cells revealed that *Acta2*, *Tagln* and *Cdh11* were already detectable at lower levels. Collectively, this suggests that mesenchymal cells lining the exocoelom have a common origin and differentiation program and are equipped to sustain rapid massive morphology changes.

A series of remarkable scRNA-seq studies have provided very valuable insight into embryo development and offered detailed reference atlases to the community ^32,56,57^. The highly useful mouse gastrulation single-cell atlas currently lacks some ExE tissues ^56^. Furthermore, identification of cell location is based solely on gene expression and can thus be confounded by similarity in transcription profiles. A RNA sequencing study of single nuclei from E9.5 to E14.5 mouse embryo placenta resolved all populations of the mature labyrinth, including fetal mesenchyme clusters characterized by *Acta2* expression ^57^. Here we provide an additional database that includes spatial and germ layer information for early ExE structures that had not been previously individualized due to absence of specific markers. Fate maps of Pre/Early Streak mouse embryos showed that amniotic ectoderm is derived from mixed fate progenitors mostly in the proximal anterior and anterolateral epiblast ^9^. Through scRNA-seq of micro-dissected amnion, we found a series of genes differentially expressed between epiblast and amnion ectoderm, including *Krt8*, *Lrp2* and *Wnt6*. *Wnt6* is a conserved amnion ectoderm marker in mouse ^58^, non-human primate ^59^, and human ^60^ embryos. *WNT6* and *LRP2* were co-expressed in a subset of human embryo ectoderm cells ^60^, similar to what we observed in amnion epithelium. As Wnt6 has been shown to regulate epithelial-mesenchymal transitions in other contexts ^61,62^, it may be compatible with an intermediate epithelial-mesenchymal state favorable for the rapid amnion expansion required upon embryo turning.

Placental defects are found in more than 50% of embryonically lethal mouse mutants ^3^. For a proportion of those genes, epiblast-specific mutants identified the germ layer responsible for lethality. *Bap1* phenotype, for example, was partially rescued by wild type trophectoderm; interrogating our atlas showed *Bap1* expression in chorion and allantois. *Crb2* phenotype however was not rescued in epiblast-specific mutants, compatible with its specific transcription in allantois. A number of mouse mutants have been shown to display defective chorio-allantoic union ^10,11^, notably through deletion of *Vcam-1* (expressed in allantois ^63^) or its partner alpha 4 integrin (*Itga4,* expressed in chorion mesoderm ^64^). Most described mutants have incomplete penetrance, suggesting there might be functional redundancy. Based on single cell profiles in allantois and chorion, we interrogated CellPhone DB ^65^ to look for ligand-receptor couples that could play a role in allantois directional growth towards chorion. To increase specificity, we focused on allantois cortex and mesenchyme versus chorion mesoderm and ectoderm (Sup5G). Significant interactions included previously described actors of chorio-allantoic fusion such as BMP2, 4, 5 and 7, FGFR2, or Wnt7b ^10,11^ as well as an array of potential new targets either within the BMP, FGF and Wnt pathways or belonging to other pathways, such as Neuropilin, Ephrin and Notch.

Although the human embryo displays a different geometry at that stage, cell populations involved in formation of amnion, placenta and umbilical cord are conserved and likely to rely on similar molecular modules. Single-cell analyses of first trimester placentas identified placental and decidual cell types at the maternal-fetal interface ^66,67^. Fibroblasts of both maternal ^66^ and fetal ^67^ origins had high *ACTA2* and *TAGLN*. Fetal fibroblasts were the primary source of *DLK1* ^67^, similar to mouse in which Dlk1 is abundant in ExE mesoderm as well as allantois and amnion mesenchyme. Time and space resolved atlases of mouse embryo and placenta are precious tools to guide interpretation of data from rare human samples.

Here we captured cytoskeleton, cell and tissue dynamics during formation of mouse embryo support organs, uncovered a major role for keratin intermediate filaments in expansion of ExE tissues, and provided a regional single cell transcriptome for mesoderm, amnion, chorion and allantois. Higher time and space resolution imaging through lattice light sheet microscopy ^68^ of K8-eYFP embryos bearing fluorescent markers for nuclei and cytoskeletal components would be very valuable to better comprehend keratins’ subcellular dynamics and interplay with actomyosin and microtubules. In order to decipher the role of KF in ExE mesenchyme at the molecular and physical level, one could take advantage of *ex vivo* explant systems for mesoderm cells ^69^ or mesoderm-derived organs such as allantois ^40,70^ from wild-type and mutant embryos. Cells could then be challenged by varying substrate rigidity and composition or knocking down potential partners within adhesion complexes. Similar studies could be undertaken on appropriate mouse and human stem cells-derived germ layer and embryo models to help decipher essential steps in the establishment of the maternal-fetal interface in human.

## ACKNOWLEDGMENTS

We wish to thank the Université Libre de Bruxelles/Erasme animal and flow cytometry facilities. We gratefully acknowledge the Université Libre de Bruxelles light microscopy (LiMiF) core facility (in particular M. Martens and J-M. Vanderwinden) for help with confocal and two-photon imaging, as well as the genomic facility for help with sequencing (A. Lefort and F. Libert). We thank Achim Gossler and Shankar Srinivas for sharing mouse lines, and Kirstie Lawson, Evangéline Despin-Guitard, Susana Chuva de Sousa Lopes, Eneko Urizar, and Joaquim Grego-Bessa for fruitful discussions.

W.N. was supported by WELBIO (SGR2015), the Université Libre de Bruxelles (ULB), and the Fonds de la Recherche Scientifique (FNRS) (PDR T.0084.16). B.S. received a FRIA fellowship of the FNRS. S.P.S. is supported by the FNRS under grant number 34772792 (MISU). I.M. is a FNRS research associate. The CMMI is supported by the European Regional Development Fund and the Walloon Region. Work in the Magin lab is supported by the Deutsche Forschungsgemeinschaft (DFG), grants MA1316/19-1 and MA1316/21-2 to T.M.M. N. S. is supported by Deutsche Forschungsgemeinschaft (DFG) grant 363055819/GRK2415. WELBIO, the FNRS, and the Fondation Erasme supported this work.

## AUTHOR CONTRIBUTIONS

W. N. and I. M. conceptualized the study, analysed and interpreted data. W.N. performed most experiments, data quantification and presentation. I. M. wrote the manuscript. S. E. E. and S. P. S participated to scRNA-seq data analysis and visualization. M. V. and D. M. performed the TEM and SEM experiments, M. V., D. M. and D.P.M. analysed and interpreted the EM data. B. S. isolated E7.25 mesoderm cells for scRNA-seq. N. S. and T. M. M. provided the K8-eYFP and *KrtyII* mouse lines, respectively, and assisted with data interpretation. A. Z. was involved in conceptualization and data interpretation and reviewed the manuscript.

## DATA AVAILABILITY

The single-cell RNA sequencing data discussed in this publication have been deposited in GEO under reference GSE167958. A shinny application is upon request. Differential gene expression is available in Embryonic GFP positive, ExE GFP positive, allantois, amnion, chorion, and combined allantois/chorion/amnion. No graph visualization means the gene was not detected in the sample.

## DECLARATION OF INTERESTS

The authors declare no conflict of interests.

## METHODS

### Mouse breeding and embryo collection

Mouse colonies were maintained in a certified animal facility in accordance with European guidelines. Experiments were approved by the local ethics committee (CEBEA). Mouse genomic DNA was isolated from ear biopsies treated for 1h at 95°C in NaOH in order to simultaneously genotype and identify animals. Mouse lines were K8-eYFP ^18^, mTmG ^71^ (Jackson laboratory), *Brachyury (T)*-Cre ^72^, *Hex*-GFP ^73^, all bred on a CD1 background, and *KrtyII* ^+/− 29^, bred on a Black6 background. Embryos were recovered at the appropriate time point after observation of a vaginal plug at day 0. Embryos were dissected in Dulbecco’s modified Eagle medium (DMEM) F-12 (Gibco) supplemented with 10% Fetal Bovine Serum (FBS), 1% Penicillin/Streptomycin and L-glutamine, and 15 mM HEPES, using #5 forceps and tungsten needles under a transmitted light stereomicroscope. Bright-field pictures of the litter or single embryo were taken before any other manipulation to ensure adequate staging. Genotyping of mutant embryos was performed after a lysis step (Lysis Buffer (Viagen)) with 1,5% Proteinase K (Qiagen).

### scRNA-seq sample preparation

*T*-Cre; mTmG embryos were collected from multiple pregnant females at E7.5, and embryos at the appropriate stage were pooled. Embryonic and ExE portions were separated by manually cutting the embryo with finely sharpened forceps. For embryonic and ExE mesoderm cells samples, MS/LS embryos were digested in dissociation buffer (1% 100 mM EDTA + 4% of 2,5% Trypsin in Phosphate Buffer saline (PBS)), and the two pure GFP positive populations were sorted through flow cytometry (FACSARIA III, BD), directly in DMEM F-12 supplemented with 10% FBS and 15 mM HEPES. For ExE structures samples, LB embryos were pooled and incubated at 4°C in pancreatic/trypsin solution (2,5% of pancreatic enzymes + 0,5% Trypsin in PBS) for 18min then washed in DMEM supplemented with 10% FBS and 1% penicillin-streptomycin. For ExE structures, allantois, amnion, and chorion were manually dissected in PBS supplemented with 0,04% BSA with finely sharpened forceps and tungsten needles. ExE structures were then incubated at 37°C in accutase (Sigma, A6964)/0,25% trypsin (Gibco, 1509-046) for 20 min (with gentle agitation at 10 min). 30% DMEM/F-12 with HEPES, 20% FBS, and 50% 4mM EDTA were added to the mix. Clumps were triturated by mouth pipetting and the solution was filtered in a non-binding Eppendorf tube with Flow MI cell 40 μm to remove debris. The solution was centrifuged at 450g for 4 min. Cells were resuspended in 50 μL in FHM (Merck, MR-024-D). For both approaches, cells were checked for viability and counted using a haemocytometer.

### scRNA-seq analysis

Single cell transcriptomics was performed with the Chromium Single Cell microfluidic device (10X Genomics). Loaded cells were individually barcoded with a 10X Chromium controller according to the manufacturer’s recommendations. The libraries were prepared using the Chromium Single Cell 3′ Library Kit (V3-chemistry, 10X Genomics), and sequenced on NovaSeq 6000 (Illumina) sequencing platform. Each sample sequenced was obtained from one experiment except for amnion that is a combination of two samples sequenced separately. 2.742 and 1.159 cells were sequenced for embryonic and ExE GFP positive cells, respectively, with a mean number of 24.619 and 56.514 reads per cell and 3.294 and 4.505 genes per cell. For the two amnion samples, 314 and 503 cells were sequenced, with a mean number of 214.164 and 133.664 reads per cell and 5.956 and 5.173 genes per cell. 3.732 and 912 cells were sequenced for chorion and allantois, respectively, with a mean number of 25.613 and 70.891 reads per cell and 3.065 and 4.747 genes per cell.

Sequencing reads were aligned and annotated with the mm10 reference dataset provided by 10X Genomics*. Cre, eGFP* and *tdTomato* sequences were added by following 10X Genomics instructions. Sequencing reads were demultiplexed using CellRanger (version 3.0.2) with default parameters. Number of genes, total counts of UMI and the percentage of mitochondrial genes were utilized for quality control. Expression value scaling and normalization, PCA and UMAP dimensionality reductions and clustering were performed using the Seurat R package (version 3.0.1)^74^. Clusters were defined using Seurat at multiple resolutions (0.2, 0.3, 0.5) and marker gene discovery was performed using the FindAllMarkers function of the Seurat package using the Wilcoxon Ranked Sum test. Markers were then selected by setting the threshold to all genes with an adjusted *p-value* lower than 0.25. Two output graphs were generated: UMAP and Dot plot. On UMAP, each point represents a cell, and its position is based on the cell embeddings determined by the PCA. Color depends on the cluster attribution. On Dot plot, the size of the dot encodes the percentage of cells expressing the gene within a cluster and the color encodes the average expression level across all cells within the cluster. Blue means high expression and grey means low expression. RNA velocity was performed on dataset processed as previously described with Seurat. Velocyto ^30^ and scvelo ^75^ packages were run on Python 3 by following the described on-line pipelines. The stochastic model was used for the RNA velocity models based on the balance of unspliced and spliced mRNA levels and their covariation. Receptor-ligand interaction analysis was performed utilizing CellPhoneDB package ^65^. As outlined in CellPhoneDB guidelines, the input data was prepared after extracting from Seurat object. Mouse gene names were converted to orthologues of human gene names which were retrieved from BioMart ^76^. The statistical method was run using the default parameters (10% threshold). Dot plot was plotted using the entire list of significant interaction pairs in the following comparisons: allantois cortex vs. chorion mesoderm, allantois mesenchyme vs. chorion mesoderm, allantois cortex vs. chorion ectoderm and allantois mesenchyme vs. chorion ectoderm. For a detailed description of the terms, please refer to the documentation of CellPhoneDB package.

### Whole-mount and section immunostaining

For immunofluorescence, embryos were fixed in PBS containing 4% paraformaldehyde for 2h at 4°C. For sagittal sections, fixed embryos were cryopreserved in 30% sucrose, embedded in OCT, and cryosectioned at 7-10 μm. Staining was performed in PBS containing 0.5% Triton X-100, 0.1% BSA, and 5% heat-inactivated horse serum. Primary antibodies were incubated overnight, and secondary antibodies were incubated for 2h at RT. Sections and whole-mount embryos were imaged on a Zeiss LSM 780 microscope equipped with Plan Apochromat 20x/0,8 M27 and an LD C Apochromat 40x/1,1 W Korr M27 objectives with 1, 3 or 5 μm Z-intervals. Image processing was done on ImageJ, Icy, or Arivis.

### Antibodies

The following primary antibodies were used: anti-Cdh1 (rabbit, 1:500, Cell Signaling, 877-616), anti-Cdh11 (rabbit, 1:200, Cell Signaling, 442S), anti-Cdx2 (rabbit, 1:200 Abcam, ab76541), anti-Cldn7 (rabbit, 1:50, Thermofisher, 34-9100), anti-Col1a1 (rabbit, 1:50, Thermofisher, PA1-85319), anti-Dlk1 (rat, 1:250, MBL, D187-3), anti-Epcam (rat, 1:100, Biolegende, 1182020), anti-Fn1 (rabbit, 1:1500, Sigma, F3648), anti-Ifitm3 (rabbit, 1:50, Abcam, ab15592), anti-GFP (goat, 1:500, Abcam, ab6673), anti-GFP (rabbit, 1:500, Life technology, A11122), anti-Kdr (rabbit, 1:100, Cell Signaling, 55B11), anti-Krt-8 (rat, 1:100, TROMA-I-S, DSHB, AB_531826), anti-LamA/C (chicken, 1:100, Novus, NBP2-25152SS), anti-LamB1 (rabbit, 1:250, Abcam, ab229025), anti-Lrp2 (rabbit, 1:50, Abcam, ab76969), anti-Nrp1 (goat, 1:50, R&D, AF566-SP), anti-Pitx1 (rabbit, 1:100, Novus, NBP1-88644), anti-Runx1 (rabbit, 1:200, Sigma, HPA004176), anti-Sox2 (rabbit, 1:100, Abcam, ab92492), anti-Stard8 (rabbit, 1:50, Biorbyt, orb101873), anti-T (rabbit, 1:100, R&D, AF2085), and anti-Vim (rabbit, 1:200, Abcam, ab92547). The following secondary antibodies were used at 1:500: anti-chicken conjugated to Alexa647 (Jackson Immunoresearch, 703-605-155), anti-goat conjugated to Alexa488 (Thermofisher, A20990) and 647 (Thermofisher A32814), anti-rabbit conjugated to Alexa488 (Thermofisher, A21206), and anti-rat conjugated to Alexa647 (Thermofisher, A21472). Alexa568 conjugated phalloidin (Invitrogen, A12380) was used at 1:200 in blocking buffer to visualize F-actin microfilaments and highlight cell membranes. Nuclei were stained with DAPI (1:1000, Sigma, D9542).

### Live Imaging

For live imaging, ectoplacental cones were conserved. Embryos were cultured in 50% DMEM-F12 with L-glutamine without phenol red, 50% rat serum (Janvier), at 37°C and 5% CO2. Embryos were observed in suspension in individual conical wells (μ-slide angiogenesis, Ibidi) to limit drift, under a Zeiss LSM 780 microscope equipped with a two-photon laser (Coherent) at 950 nm, and an LD C Apochromat 40×/ 1.1 objective. Stacks were acquired every 20 min with 5 μm Z-intervals for up to 6 h. Embryos were cultured for an additional 6-12 h after imaging to check for fitness. 3D views were processed using Arivis Vision4D v3.0 (Arivis, Germany). Registration was performed after video generation using the StackReg ImageJ plugin.

### Spectral Imaging & Image Processing

Whole-mount fixed K8-eYFP embryos were stained with DAPI for 2h in PBS containing 0.5% Triton X-100, 0.1% BSA, and 5% heat-inactivated horse serum. Spectral mode on Zeiss was used with a wavelength interval of 9 nm with a two-photon laser of 950 nm and a Z interval of 1 μm. Images were unmixed through ZEN software based on prior spectral analysis of K8-eYFP and DAPI signals alone. For big samples, the Zeiss tile option was used during acquisition and unmixed images were mounted through Huygens Professional software (Deconvolution and Stitching widget).

### Image Quantification

Quantification from confocal images was performed using the Icy software ^77^ (v2.1.0.1), ImageJ (v1.52h) and Arivis (Vision4D v3.0). Data analysis was handled through homemade Python scripts and GraphPad software (v8.4.3). Student t-tests or Mann-Whitney tests were used according to the nature of the data sets. Ns: non-significant, *p-value<0,05, **p-value<0,005, *** p-value<0,0005, and ****p-value<0,00005.

To measure the ratio of signal intensity of LamA/C over LamB1, the threshold HK means from Icy (http://icy.bioimageanalysis.org/) ^77^ on LamB1 channel was applied on several spots within specific areas. This mask was applied on the LamA/C channel. Mean intensity signals were extracted from the two channels, and the ratio LamA/C over LamB1 was calculated.

### Electron Microscopy

For Scanning Electron Microscopy, embryos were fixed with glutaraldehyde 2,5 % over night and rinsed in cacodylate buffer 0.1 M, pH 7.0. After serial dehydration in increasing concentrations of ethanol and finally acetone, samples were dried at CO_2_ critical point, then opened and mounted on Scanning Electron Microscopy stubs. Observations were performed with an ESEM Quanta F200 (FEI-ThermoFisher) microscope and secondary electron images captured with an Everhart-Thornley detector. Images were analyzed and processed by iTEM software. For Transmission Electron Microscopy analyses of thin sections, embryos were fixed with glutaraldehyde 2,5% (EM grade, Electron Microscopy Sciences) and postfixed twice in 1% OsO4 (with 1,5% ferrocyanide). Samples were stained with uranyl acetate (UA), serially dehydrated in increasing ethanol concentrations, and embedded in epoxy-resin (Agar 100 resin, Agar Scientific Ltd, UK). Sectioning was done on a Leica EM UC7 ultramicrotome and ultrathin (50-70nm thick) sections were further stained with UA and lead citrate by electron microscopy standard procedures. Observations were made on a Tecnai 10 electron microscope (FEI-ThermoFisher) and images were captured with a Veleta CCD camera and processed with SIS iTem software (Olympus).

## VIDEO LEGENDS

**Video1:** 3D reconstruction in the ExE region of a fixed whole-mount embryo stained for F-actin (Phalloidin) (n=4). Z interval is 1 μm.

**Video2**: Z-projection from two-photon live imaging of one K8-eYFP (yellow) LS embryos, when YFP becomes detectable in ExE mesoderm (n=9). Time interval is 15 min, scale bar represents 25 μm.

**Video3**: Z-projection from two-photon live imaging of a K8-eYFP (yellow) 0B embryos at 40X magnification showing whole ExE region (n=10). Time interval is 15 min, scale bar represents 25 μm.

**Video4**: Z-stack of a whole-mount fixed K8-eYFP (yellow) LB embryo stained for nuclei (DAPI, cyan) focused on ExE region (n=15). Z interval is 1 μm, scale bar represents 20 μm.

**Video5**: Z-projection from two-photon live imaging of 5 K8-eYFP (yellow) EB/LB embryos focusing on ExE region at 40X magnification highlighting K8 containing cables (n=10). Time interval is 15 min and scale bars represent 25 μm.

**Video6**: Z-projection from two-photon live imaging of 2 K8-eYFP (yellow) LB embryos focusing on ExE region at 40X magnification and showing the amnio-chorionic fold at the anterior separation point (n=6). Time interval is 15 min and scale bars represent 25 μm.

**Video7**: Z-projection from two-photon live imaging of 3 K8-eYFP (yellow) LB embryos focusing on ExE region at 40X magnification showing allantois in lateral (a, b) (n=12) and posterior (c) views (n=3). Time interval is 15 min and scale bars represent 25 μm.

**Video8 a, b**: 3D reconstructions in the ExE region of 2 fixed whole-mount K8-eYFP (yellow) embryos stained with DAPI (cyan) (n=15). Z interval is 1 μm.

---

**Supplementary Figure 1-related to Figures 1 and 2:**
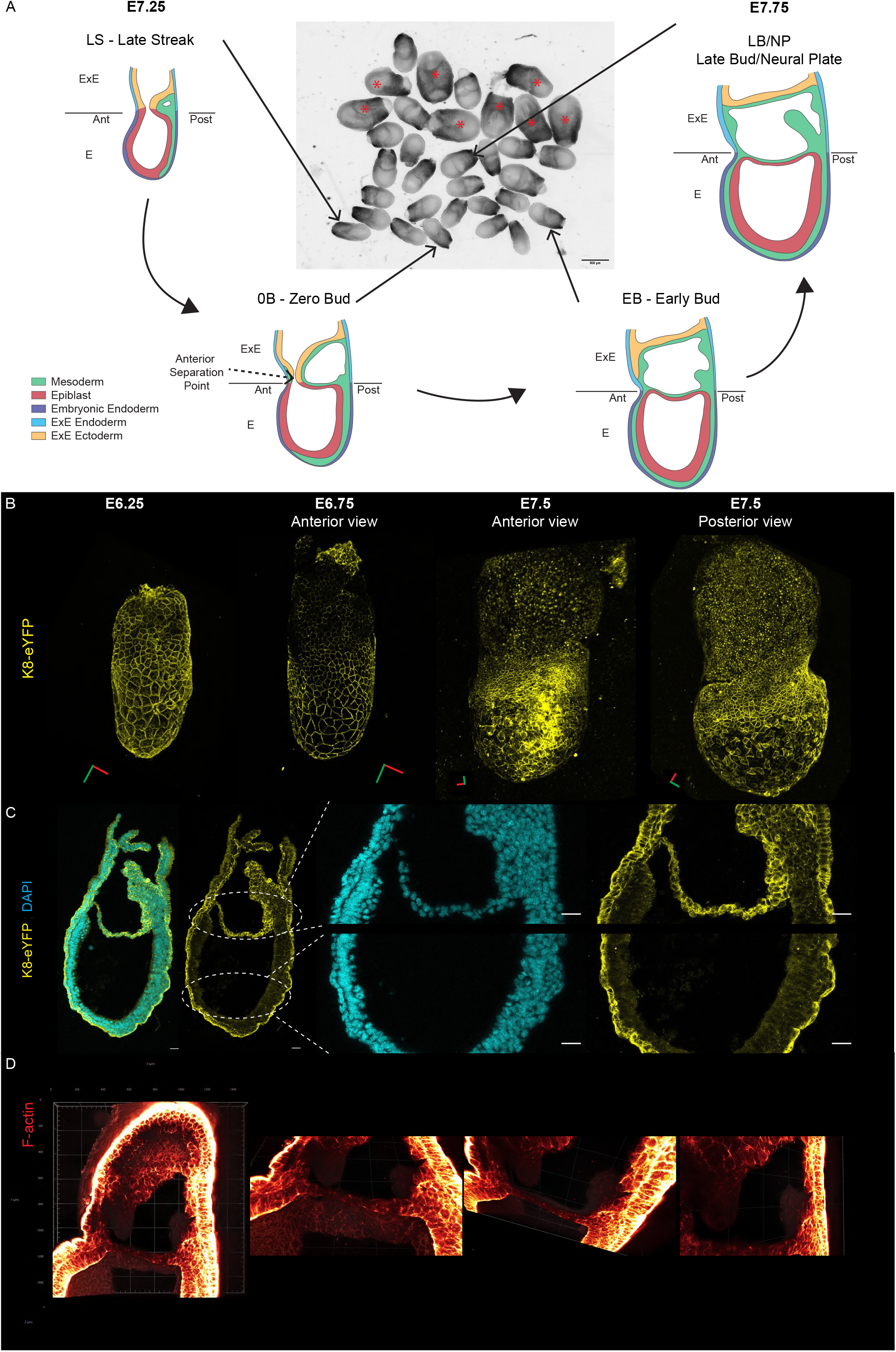
Embryo staging and cytoskeletal markers. Classification of embryos as Late Streak (LS), Zero Bud (0B), Early Bud (EB), and Late Bud/Neural Plate (LB/NP) stages based on bright-field image of two litters dissected at late E7.5. Head fold stage embryos (red stars) were not used. B. 3D reconstructions from two-photon stacks of fixed K8-eYFP embryo dissected at E6.25, E6.75, E7.5 showing visceral endoderm labelling. Green and red bars represent 40 μm in the X and Y axis, respectively. C. Z-projections of sagittal sections from a LB K8-eYFP embryo (n=30) with zooms on embryonic (Bottom) and ExE (Top) regions showing expression in endoderm as well as ExE ectoderm and mesoderm. K8-eYFP and DAPI appear in yellow and cyan. Scale bars represent 25 μm. D. 3D reconstructions of whole-mount LB embryo stained for F-actin (Phalloidin, red). Grid squares represents 200 μm.

**Supplementary Figure 2-related to Figure 5:**
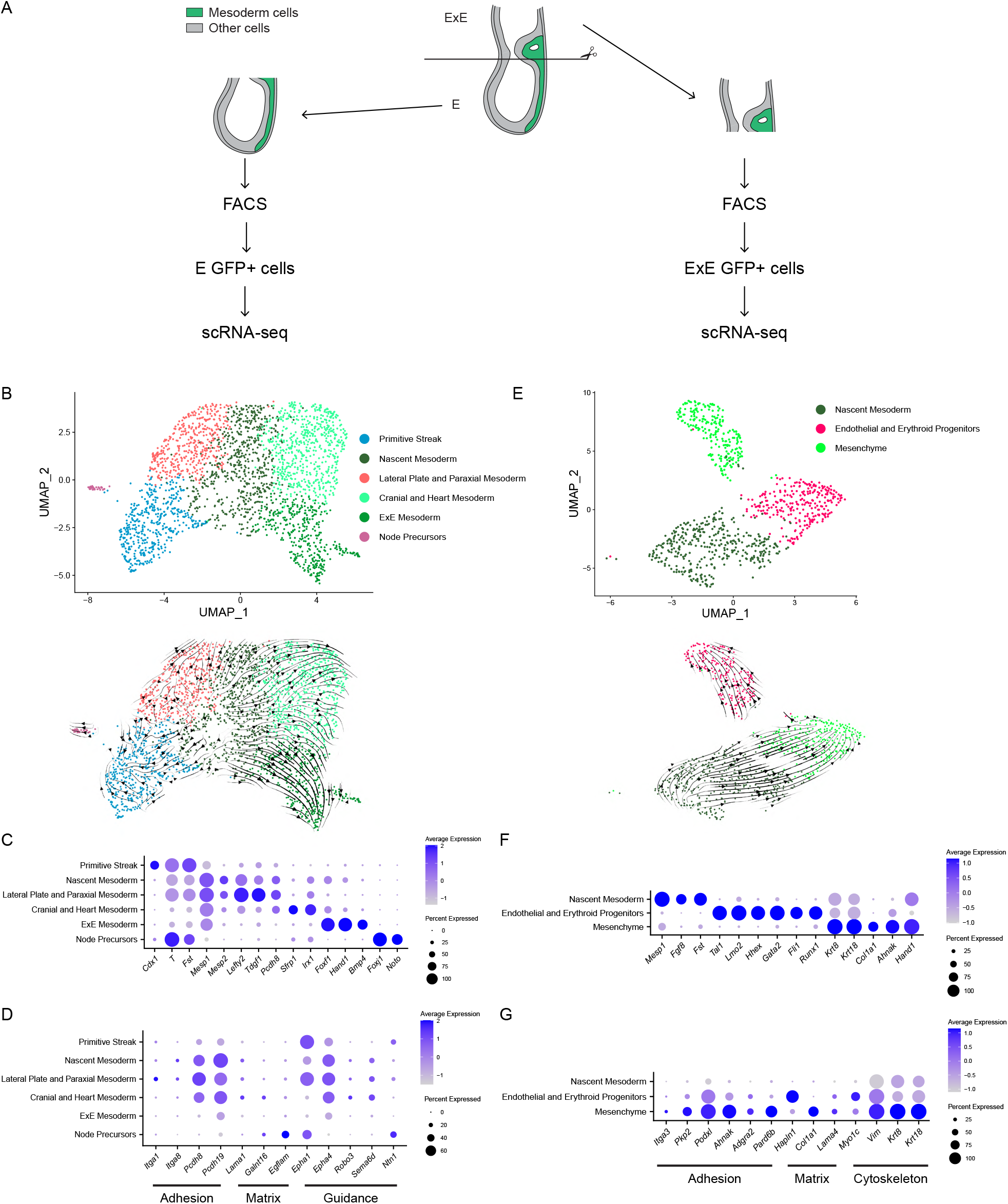
Embryonic and ExE mesoderm single cell transcriptome. A. Strategy for isolation of mesoderm cells from M/LS mTmG; T-Cre embryos. Embryos were cut manually at the embryonic/ExE border and GFP positive cells were sorted by flow cytometry. B. UMAP (Top) and RNA velocity (Bottom) of embryonic mesoderm cells where 6 unsupervised clusters were identified. C. Dot plot of specific gene expression in clusters in E region. D. Dot plot of genes previously found to have higher expression in embryonic, compared to ExE, mesoderm with roles in adhesion (Itga1, Itga8, Pcdh9, Pcdh19), matrix (Lama1, Galnt16, Egflam), and guidance (Epha1, Epha4, Robo3, Sema6d, Ntn1). E. UMAP (Top) and RNA velocity (Bottom) of ExE mesoderm cells where 3 unsupervised clusters were identified. F. Dot plot of specific gene expression in clusters in ExE region. G. Dot plot of genes previously found to have higher expression in ExE, compared to embryonic, mesoderm with roles in adhesion (Itga3, Pkp2, Podxl, Ahnak, Pard6b), matrix (Hapln1, Col1a1, Lama4), and cytoskeleton (Myoc1c, Vim, Krt8, Krt18).

**Supplementary Figure 3-related to Figure 5:**
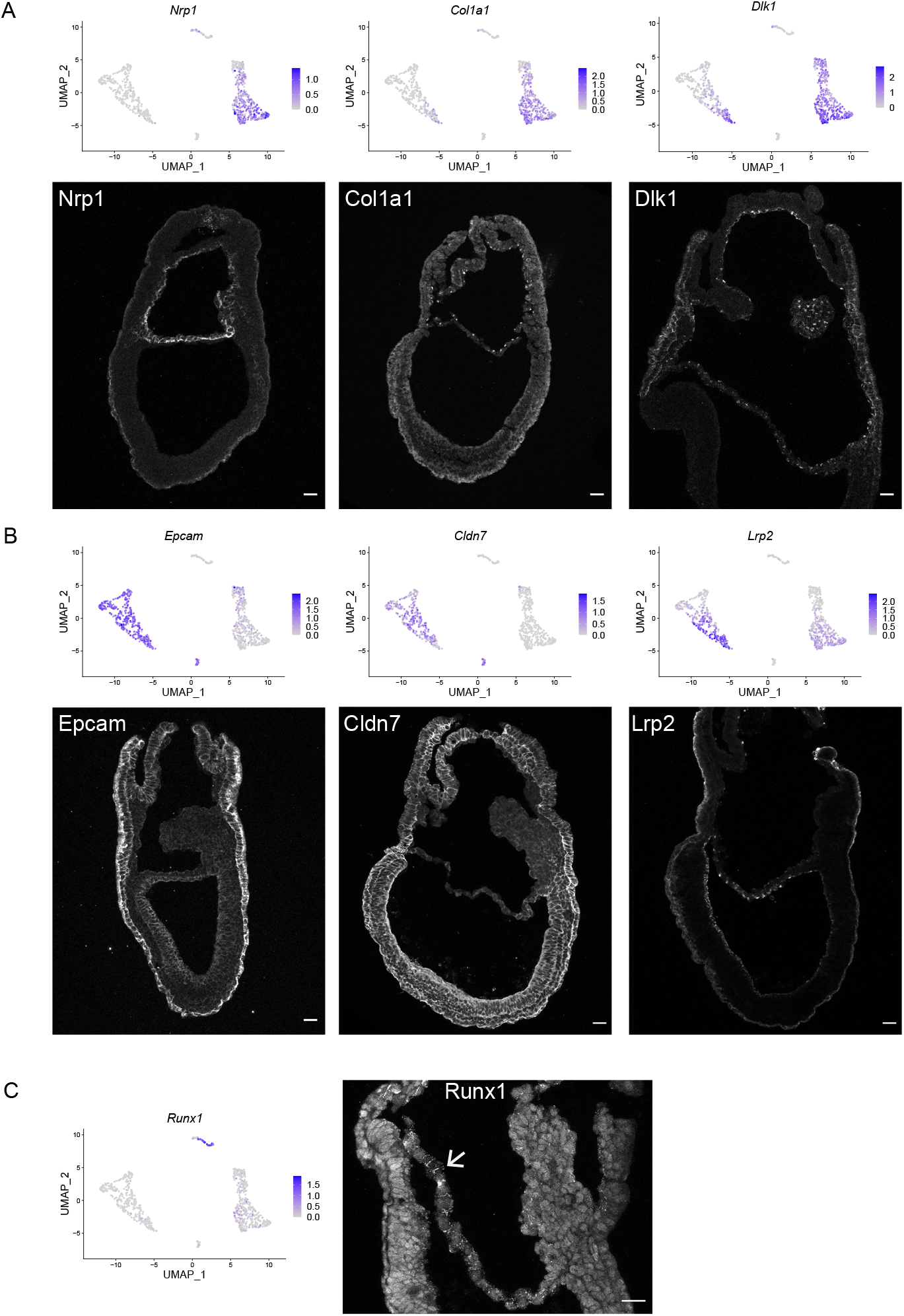
Amnion populations. A-C. UMAP and immunostaining for selected markers representing distinct clusters: Amnion Mesenchyme (A), Amnion Ectoderm (B), Erythroid Progenitors (C). Scale bars represent 25 μm.

**Supplementary Figure 4-related to Figure 6:**
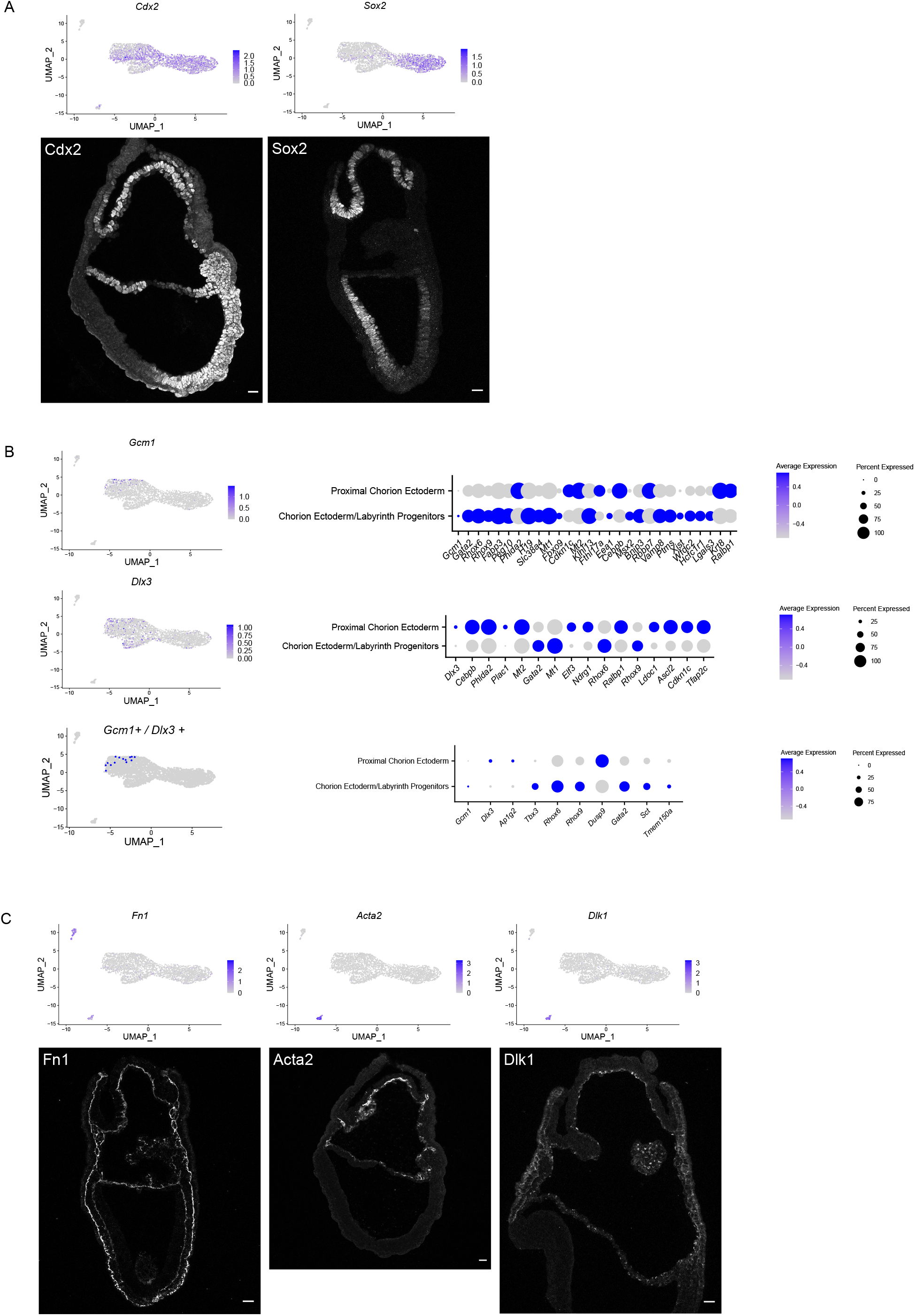
Chorion populations. A, C. UMAP and immunostaining for selected markers representing distinct clusters: Chorion Ectoderm / Labyrinth Progenitors and Trophoblasts Progenitors (A), Chorion Mesenchyme (C). B. UMAP (Left) and Dot plot of genes differentially expressed (Right) in Gcm1+ (Top, n=74), Dlx3+ (Middle, n=219), and Dlx3+/Gcm1+ (Bottom, n=14) cells.

**Supplementary Figure 5-related to Figure 7:**
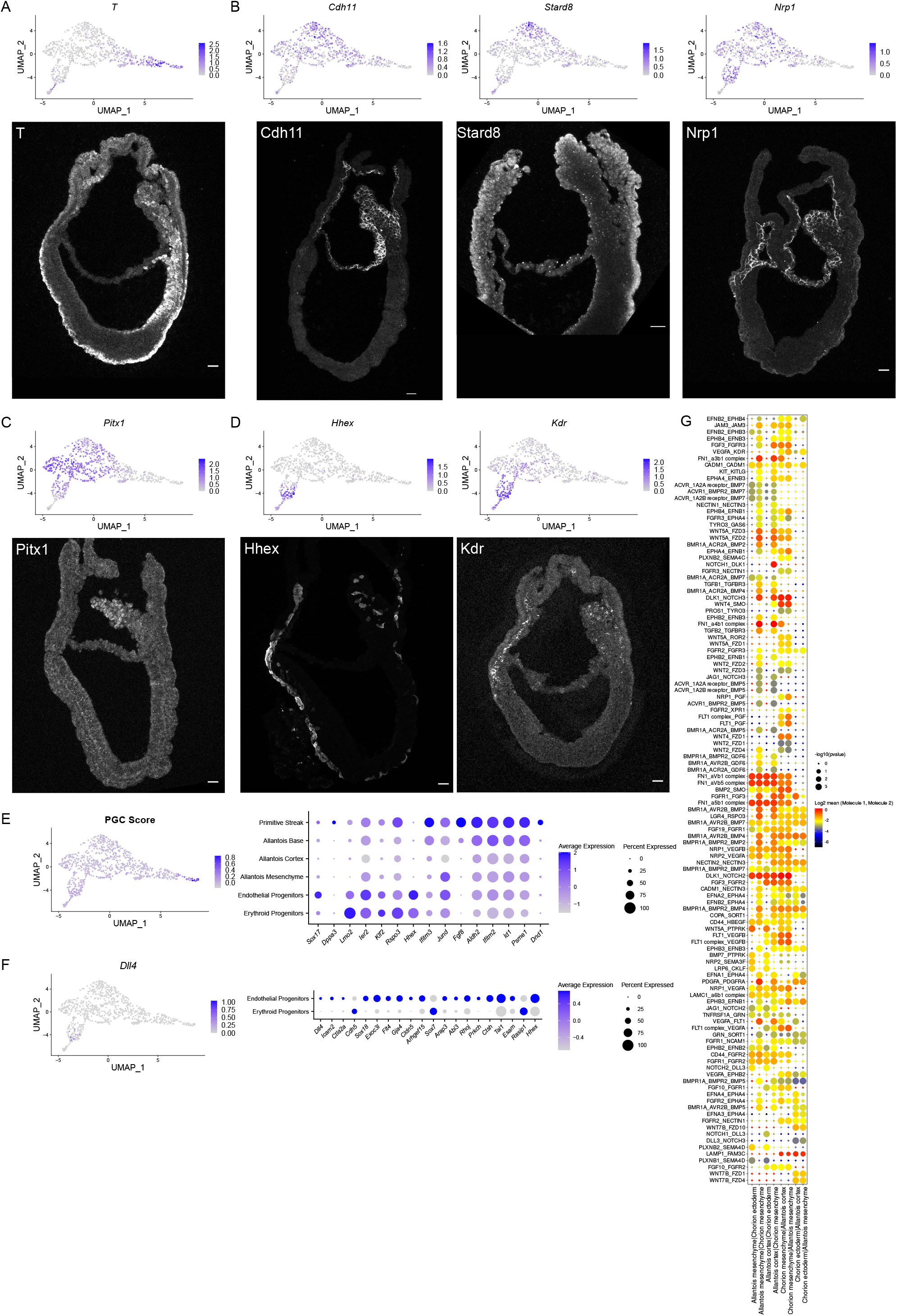
Allantois populations. A-D. UMAP and immunostaining for selected markers representing distinct clusters: Primitive Streak (A), Allantois Cortex (B), Allantois Mesenchyme (C), and Endothelial Progenitors (D). Scale bars represent 25 μm. E. UMAP of Primordial Germ Cells (PGC) score (composed of Nanog, Tfap2c, Dppa3, Sox17, Prdm1 and Nanos3) (Left) and Dot plot (Right) of genes differentially expressed in PGC. F. UMAP of Dll4 (left) and Dot plot (right) of genes differentially expressed in Dll4 positive cells among endothelial and erythroid progenitors. G. CellPhoneDB analysis revealing potential ligand-receptor pairs between chorion and allantois. Each dot’s colour intensity represents log2 of mean expression of ligand-receptor pair between the two cell types, while size of the dots indicates log of p-value. The order of interaction pair annotations is considered with respect to expression in the corresponding cell cluster. A_B pair in the vertical axis indicates interaction between clusters X|Y: the expression of partner A is considered within the first cell type (X), and the expression of partner B within the second cell type (Y). In case of more than two interaction pairs, the first two partners are considered within the first cell cluster, and the third partner within the second cell cluster.

## Notes

### Competing Interest Statement

The authors have declared no competing interest.

